# Predicting Brain Amyloid using Multivariate Morphometry Statistics, Sparse Coding, and Correntropy: Validation in 1,101 Individuals from the ADNI and OASIS Databases

**DOI:** 10.1101/2020.10.16.343137

**Authors:** Jianfeng Wu, Qunxi Dong, Jie Gui, Jie Zhang, Yi Su, Kewei Chen, Paul M. Thompson, Richard J. Caselli, Eric M. Reiman, Jieping Ye, Yalin Wang, for the Alzheimer’s Disease Neuroimaging Initiative

**Affiliations:** School of Computing, Informatics, and Decision Systems Engineering, Arizona State University, Tempe, AZ, USA; Institute of Engineering Medicine, Beijing Institute of Technology, Beijing, China; Department of Computational Medicine and Bioinformatics, University of Michigan, Ann Arbor, MI, USA; Banner Alzheimer’s Institute, 100 Washtenaw Avenue, Phoenix, AZ, USA; Imaging Genetics Center, Stevens Neuroimaging and Informatics Institute, University of Southern California, Marina del Rey, CA, USA; Department of Neurology, Mayo Clinic Arizona, Scottsdale, AZ, USA

**Author notes:** Authors contributed equally (Jianfeng Wu, Qunxi Dong). Please address correspondence to: Dr. Yalin Wang, School of Computing, Informatics, and Decision Systems Engineering, Arizona State University, P.O. Box 878809, Tempe, AZ 85287 USA, **E-mail:**. **Acknowledgments:** Data used in preparation of this article were obtained from the Alzheimer’s Disease Neuroimaging Initiative (ADNI) database (adni.loni.usc.edu). As such, the investigators within the ADNI contributed to the design and implementation of ADNI and/or provided data but most of them did not participate in analysis or writing of this report. A complete listing of ADNI investigators may be found at http://adni.loni.usc.edu/wpcontent/uploads/how_to_apply/ADNI_Acknowledgement_List.pdf.

**Keywords:** Alzheimer’s disease, Hippocampal Multivariate Morphometry Statistics (MMS), Dictionary and Correntropy-induced Sparse Coding, Beta-amyloid (Aβ) burden.

## Abstract

Biomarker-assisted preclinical/early detection and intervention in Alzheimer’s disease (AD) may be the key to therapeutic breakthroughs. One of the presymptomatic hallmarks of AD is the accumulation of beta-amyloid (Aβ) plaques in the human brain. However, current methods to detect Aβ pathology are either invasive (lumbar puncture) or quite costly and not widely available (amyloid PET). Our prior studies show that MRI-based hippocampal multivariate morphometry statistics (MMS) are an effective neurodegenerative biomarker for preclinical AD. Here we attempt to use MRI-MMS to make inferences regarding brain Aβ burden at the individual subject level. As MMS data has a larger dimension than the sample size, we propose a sparse coding algorithm, Patch Analysis-based Surface Correntropy-induced Sparse coding and max-pooling (PASCS-MP), to generate a low-dimensional representation of hippocampal morphometry for each subject. Then we apply these individual representations and a binary random forest classifier to predict brain Aβ positivity for each person. We test our method in two independent cohorts, 841 subjects from the Alzheimer’s Disease Neuroimaging Initiative (ADNI) and 260 subjects from the Open Access Series of Imaging Studies (OASIS). Experimental results suggest that our proposed PASCS-MP method and MMS can discriminate Aβ positivity in people with mild cognitive impairment (MCI) (Accuracy (ACC)=0.89 (ADNI)) and in cognitively unimpaired (CU) individuals (ACC=0.79 (ADNI) and ACC=0.81 (OASIS)). These results compare favorably relative to measures derived from traditional algorithms, including hippocampal volume and surface area, shape measures based on spherical harmonics (SPHARM), and our prior Patch Analysis-based Surface Sparse-coding and Max-Pooling (PASS-MP) methods.

## 1. INTRODUCTION

Alzheimer’s disease (AD) is a major public health concern with the number of affected individuals expected to triple, reaching 13.8 million by the year 2050 in the U.S. alone (Brookmeyer et al., 2007). Current therapeutic failures in patients with dementia due to AD may be due to interventions that are too late, or targets that are secondary effects and less relevant to disease initiation and early progression (Hyman, 2011). Preclinical AD is now viewed as a gradual process that begins many years before the onset of clinical symptoms. Measuring brain biomarkers and intervening at preclinical AD stages are believed to improve the probability of therapeutic success (Brookmeyer et al., 2007; Jack et al., 2016; Sperling et al., 2011). In the A/T/N system - a recently proposed research framework for understanding the biology of AD - the presence of abnormal levels of Aβ in the brain or cerebrospinal fluid (CSF) is used to define the presence of biological Alzheimer’s disease (Jack et al., 2016). An imbalance between production and clearance of Aβ occurs early in AD and is typically followed by the accumulation of tau protein tangles (another key pathological hallmark of AD) and neurodegeneration detectable on brain magnetic resonance imaging (MRI) scans (Hardy and Selkoe, 2002; Jack et al., 2016; Sperling et al., 2011). Brain Aβ pathology can be measured using positron emission tomography (PET) with Aβ-sensitive radiotracers, or in CSF. Even so, these invasive and expensive measurements are less attractive to subjects in preclinical stage and PET scanning is also not as widely available as MRI.

Blood-based biomarkers (BBBs) are somewhat effective for inferring Aβ burden in the brain and CSF, and are less expensive than imaging (Bateman et al., 2019; Janelidze et al., 2020; Palmqvist et al., 2020). Even so, structural MRI biomarkers are largely accessible, cost-effective, and widely used in AD imaging research as well as for clinical diagnosis. Consequently, there is great research interest in using MRI biomarkers to predict brain Aβ burden (Pekkala et al., 2020; Reisa A. Sperling et al., 2011; Tosun et al., 2016, 2014). Tosun et al. (2014) combine MRI-based measures of cortical shape and cerebral blood flow to predict Aβ status for early-MCI individuals and achieve an 83% accuracy with the LASSO approach (least absolute shrinkage and selection operator). Pekkala et al. (2020) use brain MRI measures (volumes of the cortical gray matter, hippocampus, accumbens, thalamus and putamen) to infer Aβ positivity in cognitively unimpaired (CU) subjects; they achieve a 0.70 area under the receiver operator curve (AUC) with their Disease State Index (DSI) algorithm. Although brain structural volumes are perhaps the most commonly used neuroimaging measures in AD research (Cacciaglia et al., 2018; Crivello et al., 2010; Reiter et al., 2017), surface-based subregional structure measures can offer advantages over volume measures as they contain more detailed and patient-specific shape information (Apostolova et al., 2010; Ching et al., 2020; Costafreda et al., 2011; Dong et al., 2020b, 2019; Morra et al., 2009; Qiu et al., 2009; Shen et al., 2009; Styner et al., 2004; Paul M Thompson et al., 2004; Younes et al., 2014).

Our prior studies (Shi et al., 2014; Wang et al., 2011, 2010) propose novel multivariate morphometry statistics (MMS) and apply them to analyze APOE4 dose effects on brain structures of nondemented and CU groups from the ADNI cohort (Dong et al., 2019; Li et al., 2016; Shi et al., 2014). Our proposed MMS approach uses multivariate tensor-based morphometry (mTBM) to encode morphometry along the surface tangent direction and radial distance (RD) to encode morphometry along the surface normal direction. This approach performs better for detecting clinically-relevant group differences, relative to other TBM-based methods including those using the Jacobian determinant, the largest and smallest eigenvalues of the surface metric and the pair of eigenvalues of the Jacobian matrix (Wang et al., 2011, 2010). Our recent studies (Dong et al., 2020b, 2019) show that MMS outperforms volume measures for detecting hippocampal and ventricular deformations in groups at high risk for AD at the preclinical stage. Our other related work (Wu et al., 2018) has studied hippocampal morphometry in cohorts consisting of Aβ positive AD patients (Aβ+ AD) and Aβ negative cognitively unimpaired subjects (Aβ-CU) using the MMS measure. We find significant Aβ+ AD vs. Aβ-CU group differences, using Hotelling’s *T^2^* tests. As MMS have a high dimension, it is not suitable for classification research directly. Therefore, we apply a Patch Analysis-based Surface Sparse-coding and Max-Pooling (PASS-MP) system for a low-dimensional representation of hippocampal MMS, and the binary group random forest classification of Aβ+ AD and Aβ-CU, achieving an accuracy rate of 90.48%. These studies show that MMS can distinguish clinical groups with different Aβ status. We have also successfully applied PASS-MP for MMS-based AD cognitive scores and autism spectrum disorder predictions (Dong et al., 2020a; Fu et al., 2021).

In this work, we optimize the objective function of the PASS-MP system by introducing correntropy measure (Gui et al., 2017) and propose an improved sparse coding, dubbed as the Patch Analysis-based Surface Correntropy-induced Sparse-coding and max-pooling (PASCS-MP) method. PASCS-MP does not only take the advantage of the computational efficiency of PASS-MP in its new optimization strategy, but also effectively reduces the negative influence of non-Gaussian noise in the data, which tremendously improves the prediction accuracy. PASCS-MP is an unsupervised learning method to generate a low-dimensional representation for each sample. We leverage the novel PASCS-MP method on MMS to further explore hippocampal morphometry differences for the following contrasts at the individual subject level: (1) Aβ positive individuals with mild cognitive impairment (Aβ+ MCI) vs. Aβ negative individuals with mild cognitive impairment (Aβ-MCI) from ADNI, and (2) Aβ positive cognitively unimpaired subjects (Aβ+ CU from ADNI and OASIS) versus Aβ negative cognitively unimpaired subjects (Aβ-CU from ADNI and OASIS). We apply the proposed PASCS-MP and a binary random forest classifier to classify individuals with different Aβ status. We hypothesize that our MMS-based PASCS-MP may provide stronger statistical power relative to traditional hippocampal volume, surface area and spherical harmonics (SPHARM) based hippocampal shape measurements, in predicting subjects’ Aβ status. We expect that the knowledge gained from this type of research will enrich our understanding of the relationship between hippocampal atrophy and AD pathology, and thus help in assessing disease burden, progression, and treatment effects.

## 2. SUBJECTS and METHODS

### 2.1 Subjects

Data for testing the performance of our proposed framework and comparable methods are obtained from the ADNI database (Mueller et al., 2005, adni.loni.usc.edu) and the OASIS database (Marcus et al., 2010). ADNI is the result of efforts of many co-investigators from a broad range of academic institutions and private corporations. Subjects are recruited from over 50 sites across the U.S. and Canada. The primary goal of ADNI is to test whether biological markers, such as serial MRI and positron emission tomography (PET), combined with clinical and neuropsychological assessments, can measure the progression of MCI and early AD. Subjects originally recruited for ADNI-1 and ADNI-GO have the option to be followed in ADNI-2. For up-to-date information, see www.adniinfo.org.

From the ADNI cohort, we analyze 841 age and sex-matched subjects with florbetapir PET data and T1-weighted MR images, including 151 AD patients, 342 MCI and 348 asymptomatic CU individuals. Among them, all the 151 AD patients, 171 people with MCI and 116 CU individuals were Aβ positive. The remaining 171 MCI and 232 CU individuals were Aβ negative. From OASIS database, we analyze age-and-sex-matched 260 subjects with florbetapir PET data and T1-weighted MR images, including 52 Aβ positive CU and 208 Aβ negative CU.

In addition to each MRI scan, we also analyze the corresponding Mini-Mental State Exam (MMSE) scores (Folstein et al., 1975) and centiloid measures (Navitsky et al., 2018). Operationally, the *positivity* of Aβ biomarkers is defined using standard cut-offs, with some efforts to reconcile differences among different Aβ radiotracers using a norming approach called the centiloid scale (Klunk et al., 2015; Rowe et al., 2017). ADNI florbetapir PET data are processed using AVID pipeline (Navitsky et al., 2018), and OASIS florbetapir PET data are processed using PUP (Lee et al., 2013; Su et al., 2015). Both are converted to the centiloid scales according to their respective conversion equations (Navitsky et al., 2018; Su et al., 2019). A centiloid cutoff of 37.1 is used to determine Aβ positivity, this threshold corresponds to pathologically determined moderate to frequent plaques (Fleisher et al., 2011). **Table 1** shows demographic information we analyze from the ADNI and OASIS cohorts.

**Table 1.** Demographic information for the subjects we study from the ADNI and OASIS cohorts.

| Database | Group | Sex (M/F) | Age | MMSE | Centiloid |
| --- | --- | --- | --- | --- | --- |
| ADNI Cohort | Aβ+ AD (n=151) | 79/72 | 74.6±7.8 | 22.6±3.1 | 86.3±27.4 |
| | Aβ+ MCI (n=171) | 92/79 | 74.1±7.4 | 27.7±1.7 | 76.8±26.4 |
| | Aβ- MCI (n=171) | 92/79 | 74.0±7.4 | 28.3±1.6 | 8.9±14.9 |
| | Aβ+ CU (n=116) | 45/71 | 75.9±6.1 | 28.9±1.1 | 71.1±26.4 |
| | Aβ- CU (n=232) | 90/142 | 75.7±6.3 | 29.0±1.3 | 7.5±14.5 |
| OASIS Cohort | Aβ+ CU (n=52) | 22/30 | 70.5±7.5 | 29.0±1.3 | 71.4±20.9 |
| | Aβ- CU (n=208) | 88/120 | 68.5±6.8 | 29.0±1.3 | 8.5±9.5 |

Values are mean ± standard deviation where applicable.

### 2.2 Proposed pipeline

This work develops the PASCS-MP framework to predict individual Aβ burden (see **Fig. 1** for the processing pipeline). In panel (1), hippocampal structures are segmented from registered brain MR images with FIRST from the FMRIB Software Library (FSL) (Paquette et al., 2017; Patenaude et al., 2011). Hippocampal surface meshes are constructed with the marching cubes algorithm (Lorensen and Cline, 1987). In panel (2), hippocampal surfaces are parameterized with the holomorphic flow segmentation method (Wang et al., 2007). After the surface fluid registration algorithm, the hippocampal MMS features are calculated at each surface point. We propose a PASCS-MP and classification system to refine and classify MMS patches in individuals with different Aβ status. We randomly select patches on each hippocampal surface and generate a sparse code for each patch with our novel PASCS. Next, we adopt a max-pooling operation on the learned sparse codes of these patches to generate a new representation (a vector) for each subject. Finally, we train binary random forest classifiers on individual sparse codes in people with different Aβ status; we validate them with 10-fold cross-validation. The whole system is publicly available^1^.

**Fig. 1.**
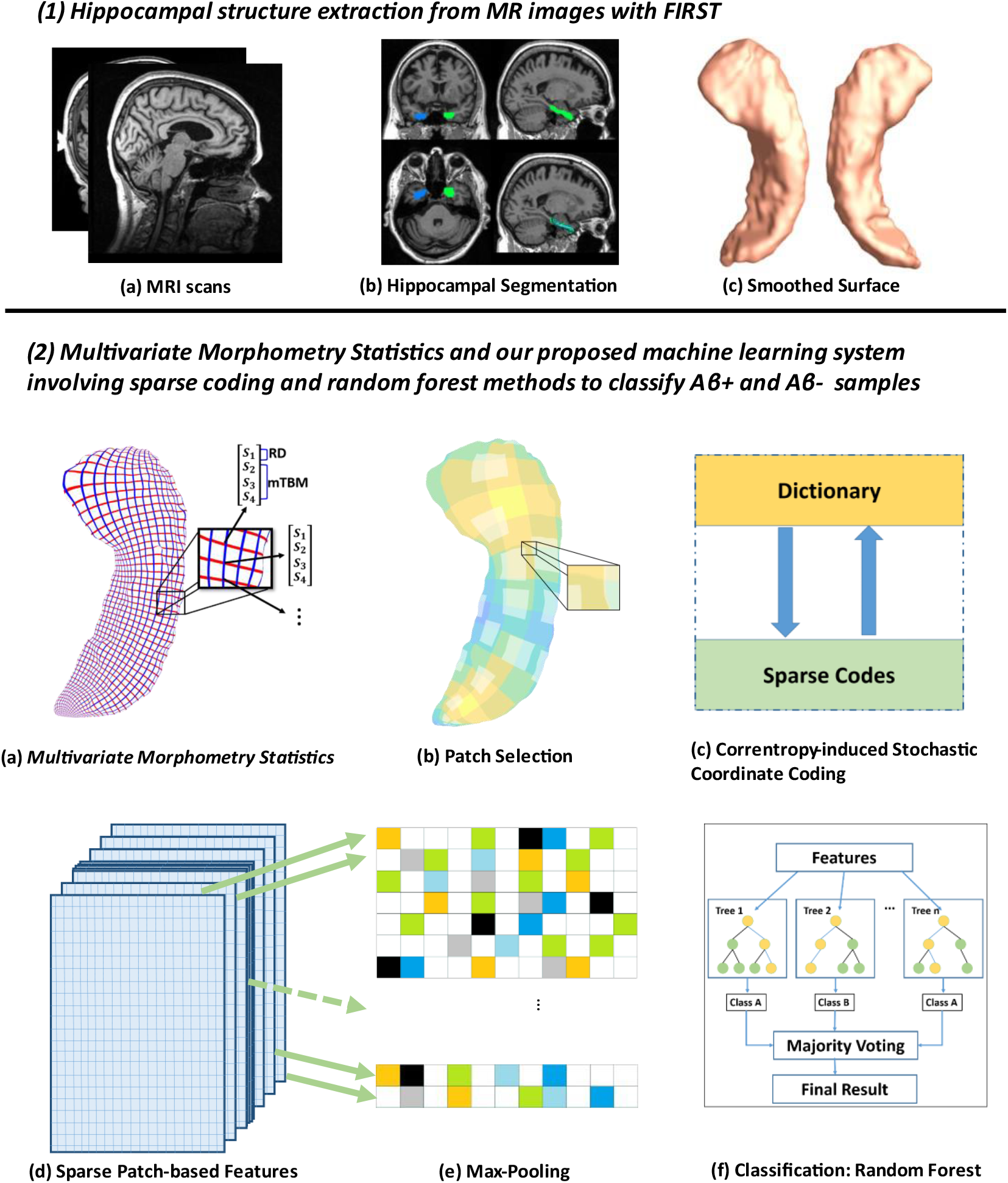
System pipeline. Panel (1) shows hippocampal surfaces generated from brain MRI scans. In panel (2), surface-based Multivariate Morphometry Statistics (MMS) are calculated after fluid registration of surface coordinates across subjects. MMS is a 4 × 1 vector on each vertex, including radial distance (scalar) and multivariate tensor-based morphometry (3 × 1 vector). We randomly select patches on each hippocampal surface and generate a sparse code for each patch with our novel Patch Analysis-based Surface Correntropy-induced Sparse-coding (PASCS) method. Next, we apply the max pooling operation to the learned sparse codes to generate a new representation (a vector) for each subject. Finally, we train binary random forest classifiers on these representations and validate them with 10-fold cross-validation.

#### 2.2.1 Image Processing

Firstly, we use FIRST (FMRIB’s Integrated Registration and Segmentation Tool) (Patenaude et al., 2011) to segment the original MRI data and map the hippocampus substructure. After obtaining a binary segmentation of the hippocampus, we use a topology-preserving level set method (Han et al., 2003) to build surface models. Based on that, the marching cubes algorithm (Lorensen and Cline, 1987) is applied to construct triangular surface meshes. Then, to reduce the noise from MR image scanning and to overcome partial volume effects, surface smoothing is applied consistently to all surfaces. Our surface smoothing process consists of mesh simplification using *progressive meshes* (Hoppe, 1996) and mesh refinement by the Loop subdivision surface method (Loop, 1987). Similar procedures adopted in a number of our prior studies (Colom et al., 2013; Luders et al., 2013; Monje et al., 2013; Shi et al., 2015, 2013b, 2013a; Wang et al., 2012, 2010) have shown that the smoothed meshes are accurate approximations to the original surfaces, with a higher signal-to-noise ratio (SNR).

To facilitate hippocampal shape analysis, we generate a conformal grid (150 × 100) on each surface, which is used as a canonical space for surface registration. On each hippocampal surface, we compute its conformal grid with a holomorphic 1-form basis (Wang et al., 2010; Wang et al., 2007). We adopt surface conformal representation (Shi et al., 2015, 2013a) to obtain surface geometric features for automatic surface registration. This consists of the conformal factor and mean curvature, encoding both intrinsic surface structure and information on its 3D embedding. After we compute these two local features at each surface point, we compute their summation and then linearly scale the dynamic range of the summation into the range 0-255, to obtain a feature image for the surface. We further register each hippocampal surface to a common template surface. With surface conformal parameterization and conformal representation, we generalize the well-studied image fluid registration algorithm (Bro-Nielsen and Gramkow, 1996; Agostino et al., 2003) to general surfaces. Furthermore, most of the image registration algorithms in the literature are not symmetric, i.e., the correspondences between the two images depending on which image is assigned as the deforming image and which is the non-deforming target image. An asymmetric algorithm can be problematic as it tends to penalize the expansion of image regions more than shrinkage (Rey et al., 2002). Thus, in our system, we further extend the surface fluid registration method to an inverse-consistent framework (Leow et al., 2005). The obtained surface registration is diffeomorphic. For details of our inverse-consistent surface fluid registration method, we refer to (Shi et al., 2013a).

#### 2.2.2 Surface-based Morphometry Feature Extraction

After parameterization and registration, we establish a one-to-one correspondence map between hippocampal surfaces. This makes it effective for us to compare and analyze surface data. Besides, each surface has the same number of vertices (150 × 100) as shown in panel 2 of **Fig. 1**. The intersection of the red curve and the blue curve is a surface vertex, and at each vertex, we adopt two features, the radial distance (RD) and the surface metric tensor used in multivariate tensor-based morphometry (mTBM). The RD (a scalar at each vertex) represents the thickness of the shape at each vertex to the medical axis (Pizer et al., 1999; Thompson et al., 2004), this reflects the surface differences along the surface normal directions. The medial axis is determined by the geometric center of the isoparametric curve on the computed conformal grid (Wang et al., 2011). The axis is perpendicular to the isoparametric curve, so the thickness can be easily calculated as the Euclidean distance between the core and the vertex on the curve. The mTBM statistics (a 3 × 1 vector at each vertex) have been frequently studied in our prior work (Shi et al., 2015, 2013b; Wang et al., 2010, 2009). They measure local surface deformation along the surface tangent plane and show improved signal detection sensitivity relative to more standard tensor-based morphometry (TBM) measures computed as the determinant of the Jacobian matrix (Wang et al., 2013). RD and mTBM jointly form a new feature, known as the surface multivariate morphometry statistics (MMS). Therefore, MMS is a 4 × 1 vector at each vertex. The surface of the hippocampus in each brain hemisphere has 15,000 vertices, so the feature dimensionality for each hippocampus in each subject is 60,000.

#### 2.2.3 Surface Feature Dimensionality Reduction

The above mentioned vertex-wise surface morphometry feature, MMS, is a high-fidelity measure to describe the local deformation of the surface and can provide detailed localization and visualization of regional atrophy or expansion (Yao et al., 2018) and development (Thompson et al., 2000). However, the high dimensionality of such features is likely to cause problems for classification. Feature reduction methods proposed by (Davatzikos et al., 2008; Sun et al., 2009) may ignore the intrinsic properties of a structure’s regional morphometry. Therefore, we introduce the following feature reduction method for the vertex-wise surface morphometry features.

The surface MMS feature dimension is typically much larger than the number of subjects, i.e., the so-called *high dimension-small sample problem*. To extract useful surface features and reduce the dimension before making predictions, this work first randomly generates square windows on each surface to obtain a collection of small image patches with different amounts of overlap. In our prior AD studies (Wu et al., 2018; Zhang et al., 2016a, 2016b), we discuss the most suitable patch size and number. Therefore, in this work, we adopt the same optimal experimental settings, as 1,008 patches (patch size=10 × 10 vertices) for each subject (504 patches for each side of the hippocampal surface). As these patches are allowed to overlap, a vertex may be contained in several patches. The zoomed-in window in subfigure (b) of panel in **Fig.1** shows overlapping areas on selected patches. After that, we use the technique of sparse coding and dictionary learning (Mairal et al., 2009) to learn meaningful features. Dictionary learning has been successful in many image processing tasks as it can concisely model natural image patches. In this work, we propose a novel sparse coding and dictionary learning method with an *l1*-regularized correntropy loss function named *Correntropy-induced Sparse-coding (CS)*, which is expected to improve the computational efficiency compared to Stochastic Coordinate Coding (SCC) (Lin et al., 2014). Formally speaking, correntropy is a generalized similarity measure between two scalar random variables U and V, which is defined by 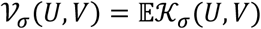. Here, 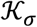 is a Gaussian kernel given by 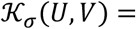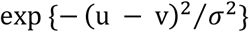 with the scale parameter σ > 0, (u-v) being a realization of (U, V) (Feng et al., 2015; Gui et al., 2017). Utilizing the correntropy measure as a loss function will reduce the negative influence of non-Gaussian noise in the data.

Classical dictionary learning techniques (Lee et al., 2007; Olshausen and Field, 1997) consider a finite training set of feature maps, *X* = (*x*_1_, *x*_2_, …, *x*_*n*_) in *R*^*p*×*n*^. In our study, *X* is the set of MMS features from *n* surface patches of all the samples. All the MMS features on each surface patch, *x*_*i*_, is reshaped to a *p*-dimensional vector. And we desire to generate a new set of sparse codes, *Z* = (*z*_1_, *z*_2_, …, *z*_*n*_) in *R*^*m*×*n*^ for these features. Therefore, we aim to optimize the empirical cost function as **Eq. (1)**.

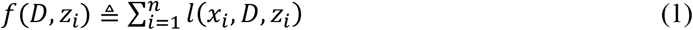

where *D* ∈ *R*^*p*×*m*^ is the dictionary and *z*_*i*_ ∈ *R*^*m*^ is the sparse code of each feature vector. *l*(*x*_*i*_, *D*, *z*_*i*_) is the loss function that measures how well the dictionary *D* and the sparse code *z*_*i*_ can represent the feature vector *x*_*i*_. Then, *x*_*i*_ can be approximated by *x*_*i*_ = *Dz*_*i*_. In this way, we convert the *p*-dimensional feature vector, *x*_*i*_, to a *m*-dimensional sparse code, *z*_*i*_, where *m* is the dimensionality of the sparse code and the dimensionality could be arbitrary. In this work, we introduce the correntropy measure (Gui et al., 2017) to the loss function and define the *l*_1_-sparse coding optimization problem as **Eq. (2)**

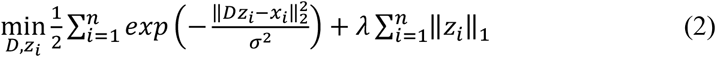

where *λ* is the regularization parameter, σ is the kernel size that controls all properties of correntropy. ‖∙‖_2_ and ‖∙‖_1_are the *l*_2_-norm and *l*_1_-norm and *exp()* represents the exponential function. The first part of the loss function measures the degree of the image patches’ goodness and the correntropy may help remove outliers. Meanwhile, the second part is well known as the *l*_1_ penalty (Fu, 1998) that can yield a sparse solution for *z*_*i*_ and select robust and informative features. Specifically, there are *m* columns (atoms) in the dictionary *D* and each atom is *d*_*j*_ ∈ *R*^*p*^, *j* = 1, 2, …, *m*. To avoid *D* from being arbitrarily large and leading to arbitrary scaling of the sparse codes, we constrain each *l*_2_-norm of each atom in the dictionary no larger than one. We will let *C* become the convex set of matrices verifying the constraint as **Eq. (3)**.

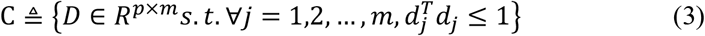

Note that, the empirical problem cost *f*(*D*, *z*_*i*_) is not convex when we jointly consider the dictionary *D* and the coefficients *Z*. But the function is convex concerning each of the two variables, *D*, and *Z*, when the other one is fixed. Since it takes much time to solve *D* and *Z* when dealing with large-scale data sets and a large-size dictionary, we adopt the framework in the stochastic coordinate coding (SCC) algorithm (Lin et al., 2014), which can dramatically reduce the computational cost of the sparse coding, while keeping a comparable performance.

To solve this optimization problem, we reformulate the first part of the equation by the half-quadratic technique (Nikolova and Ng, 2006) and then the objective can be solved as the minimization problem **Eq.(4)**:

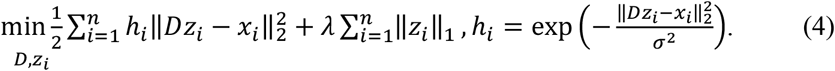

Here the auxiliary variable, ℎ_*i*_, will be updated in each update iteration. At each iteration, we update *D* and *Z* alternately, which means we firstly fix *D* and update the sparse code *Z* with coordinated descent (CD) and then fix Z to update the dictionary *D* via stochastic gradient descent (SGD).

As our optimization method is stochastic, we only update the sparse code and dictionary with only one signal for each iteration. In the following paragraphs, we will discuss the optimization in one iteration with only one signal. If a signal, *x* = (*x*_1_, *x*_2_, …, *x*_*p*_) ^*T*^ ∈ *R*^*p*^, is given, we first update its corresponding sparse code, *z* = (*z*_1_, *z*_2_, …, *z*_*m*_), via CD. Let *z*_*l*_ denote the *l* -th entry of *z* and *d*_*kl*_ represents the *k* -th item of *d*_*l*_ . *d*_*l*_ is the *l* -th atom/column of the dictionary *D*. Then, we can calculate the partial derivative of *z*_*l*_ in the first part of the function, *f*(*D*, *z*_*i*_), as **Eq. (5)**

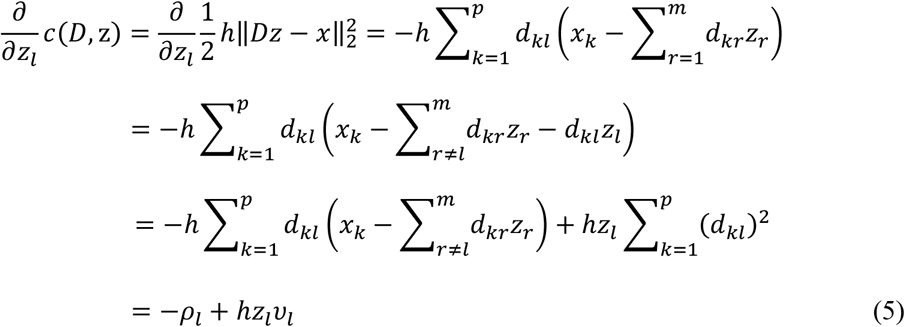

Where 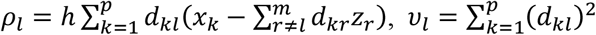 and *h* is the auxiliary variable for the signal. Since we normalize the atom, *d*_*l*_, in each iteration, *v*_*l*_ can be ignored. Then, we compute the subdifferential of the lasso loss function and equate it to zero to find the optimal solution as follows:

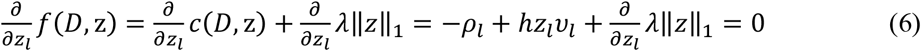

Then, according to the derivative of the *l_1_*-norm, we can have the following equations.

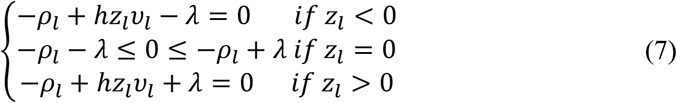

Finally, we can get the soft thresholding function as:

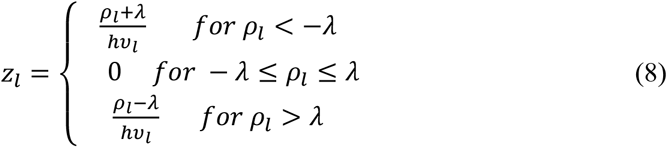

After we update the sparse code, we propose the following strategy to accelerate the convergence for updating the dictionary *D*. The atom, *d*_*l*_ will stay unchanged if *z*_*l*_ is zero since 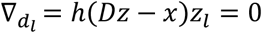. Otherwise, as shown in **Fig. 2**, we can update the *l*-th atom of the dictionary *D* as *d*_*l*_ ← *d*_*l*_ − γ_*l*_*h*(*Dz* − *x*)*z*_*l*_ . γ_*l*_ is the learning rate provided by an approximation of the Hessian: *R* ← *R* + *zz*^*T*^ and γ_*l*_ is given by 1/*r*_*ll*_, where *r*_*ll*_ is the item at the *l*-th row and *l*-th column of the Hessian matrix *R*. The pseudo-code of the model was shown in **Alg. 1**, dubbed as PASCS.

**Fig. 2.**
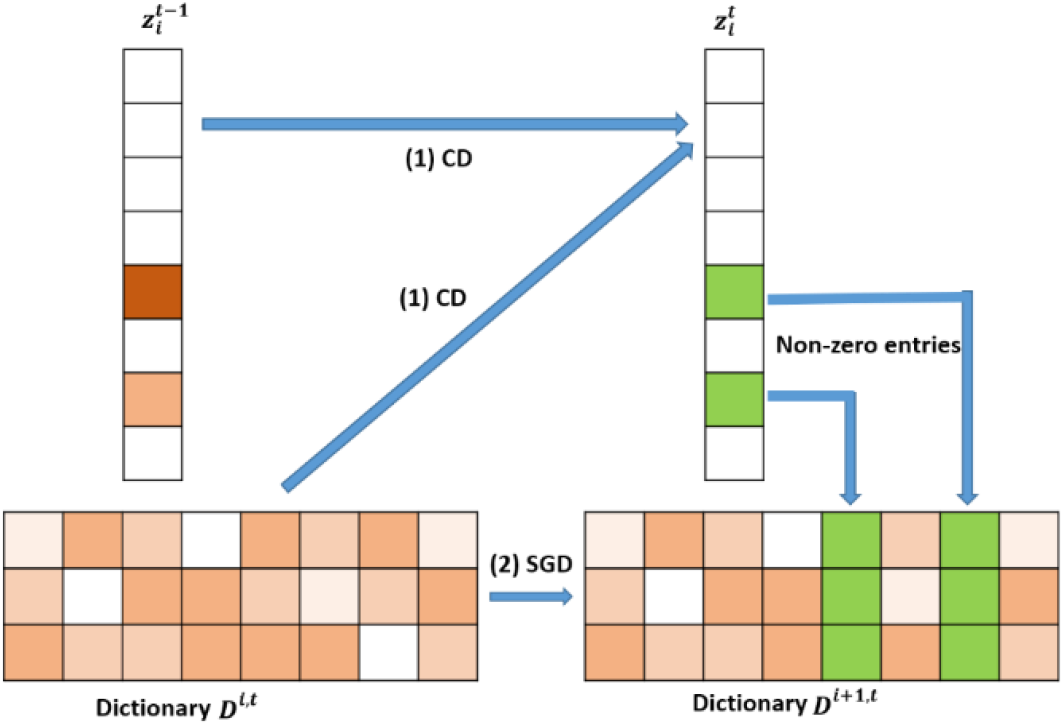
Illustration of one iteration of the proposed Patch Analysis-based Surface Correntropy-induced Sparse-coding (PASCS) algorithm. The input is many 10 × 10 patches on each surface based on our multivariate morphometry statistics (MMS). With an image patch *xi*, PASCS performs one step of coordinate descent (CD) to find the support and the sparse code. Meanwhile, PASCS performs a few steps of CD on supports (non-zero entries) to obtain a new sparse code 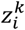. Then, PASCS updates the supports (*green boxes in the figure*) of the dictionary by stochastic gradient descent (SGD) to obtain a new dictionary *D*^*i*+1,*t*^. Here*, t* represents the *t-*th epoch; *i* represents the *i*-th patch.

**Alg. 1.**
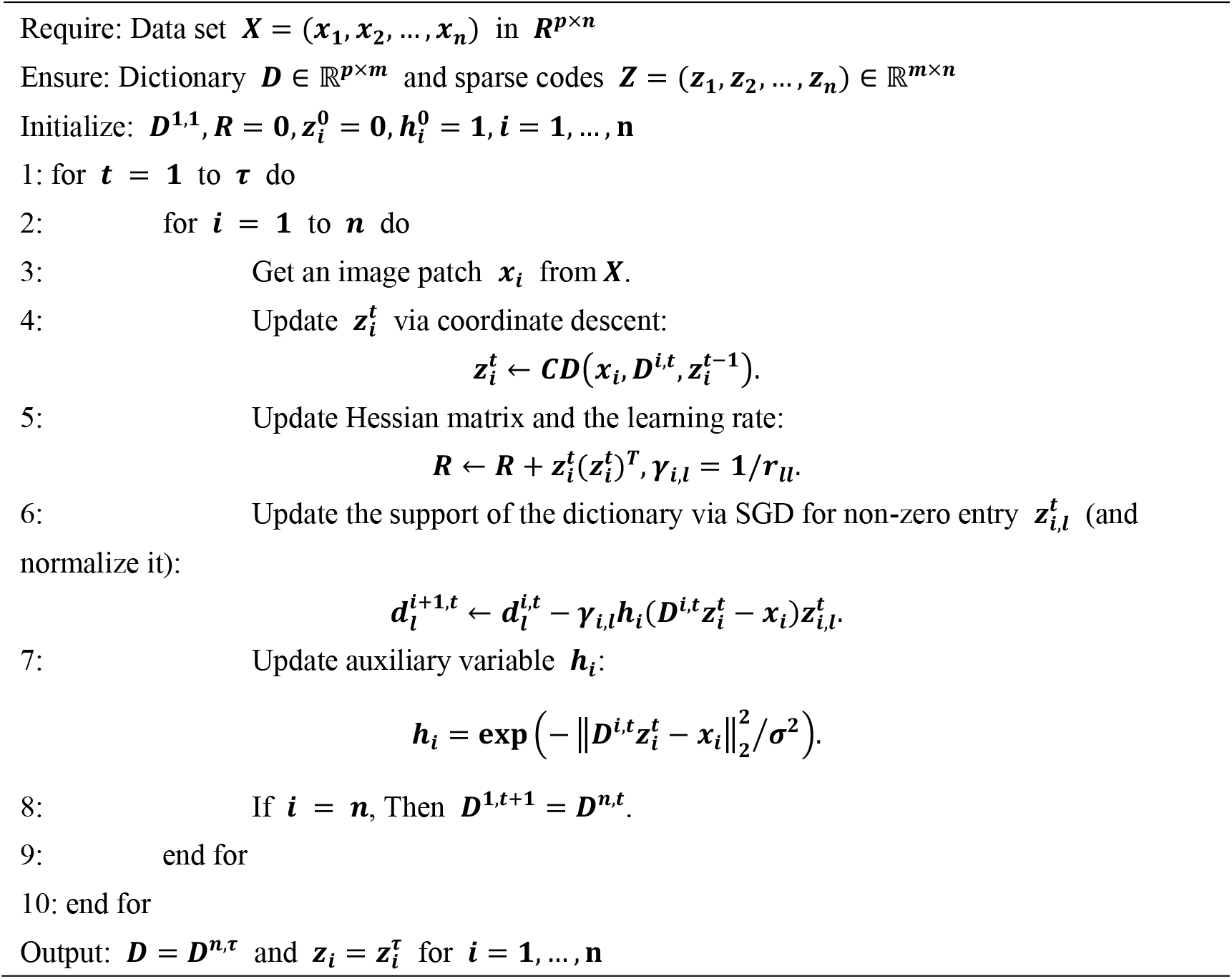
Patch Analysis-based Surface Correntropy-induced Sparse-coding.

#### 2.2.4 Pooling and Classification

After we get the sparse code (the dimension is *m*) for each patch, the dimensionality of sparse codes for each subject is still too large for classification, which is *m* × 1008. Therefore, we apply Max-pooling to reduce the feature dimensionality for each subject. Max-pooling (Boureau et al., 2010) is a way of taking the most responsive node of a given region of interest and serves as an important layer in the convolutional neural network architecture. In this work, we compute the maximum value of a particular feature over all sparse codes of a subject and generate a new representation for each subject, which is an *m*-dimensional vector. These summary representations are much lower in dimension, compared to using all the extracted surface patch features; this can improve results generalizability via less over-fitting.

With these dimension-reduced features, we choose the random forest algorithm (Liaw and Wiener, 2002) for the binary classification. Random forests are a combination of tree predictors such that each tree depends on the values of a random vector sampled independently and with the same distribution for all trees in the forest. This algorithm adopts a learning process called *feature bagging*. In this process, we select a random subset of the features several times and then train a decision tree for each subset. If some features are strong predictors of the response, they will be selected in many decision trees and this makes them correlated. In comparison with decision trees, random forests have the same bias but lower variance, which means they can overcome the drawback of overfitting caused by a small data set. For our sparse surface features, when the size of the training set becomes small, diversification becomes more subtle, and the method can better detect these subtle differences. In this project, we use the random forest classifier in the *scikit-learn* package (https://scikit-learn.org/) with the default settings. Besides, under the imbalanced-data condition (such as 116 Aβ+ CU and 232 Aβ-CU in the ADNI data set), the classifier tends to classify all the training data into the major class, as it aims to maximize training accuracy. Therefore, we adopt *random undersampling* (Dubey et al., 2014) to balance the numbers of training subjects in the two classes. All the experiments in this work use the same setups for the random forest classifier and random undersampling.

### 2.3 Performance Evaluation Protocol

Before using hippocampal MMS features for Aβ status classification, we need to apply PASCS-MP to extract sparse codes from these high dimensional MMS features. The performance of PASCS-MP has a close relationship to four key parameters: the patch size, the dimensionality of the learned sparse coding, the regularization parameter for the *l*_1_-norm (*λ*), and the kernel size (σ) in the exponential function (see **Eq.(2)**). Patch-based analysis has been widely used for image segmentation and classification (Kao et al., 2020). Leveraging patches in our MMS can preserve well the properties of the regional morphometry of the hippocampal surface since the vertices that carry strong classification power are always clustered on the surface and a set of such vertices typically has a stronger classification ability compared to using just a single vertex. However, the size of the set of such vertices is unknown. Therefore, we select the vertices by randomly selecting the same number of square patches with different sizes and compared the performance of the final classification accuracy for the different patch sizes. The dimensionality of the learned sparse coding (*m*) is also the dimensionality of the representation for each subject. The model might miss some significant information if the dimensionality is too low. Also, the representations will contain too much redundant information when the dimensionality is too large. The regularization parameter for the *l1*-norm (λ) will control the sparsity of the learned sparse codes. A suitable regularization parameter will select significant features meanwhile reducing noise. The kernel size in the exponential function controls all properties of correntropy. Correntropy is directly related to the probability of how similar two random variables are in a neighborhood of the joint space controlled by the kernel bandwidth, i.e., the kernel bandwidth acts as a zoom lens, controlling the *observation window* over which similarity is assessed. This adjustable window provides an effective mechanism to eliminate the detrimental effect of outliers (Liu et al., 2007).

Thus, we adopt 10-fold cross-validation to evaluate the classification accuracy on another dataset from ADNI 2 with a series of key parameter candidates and select the optimal parameter setups. The detailed information about the dataset and the key parameter candidates will be introduced in next section. For the 10-fold cross-validation, we randomly shuffle and split the dataset into ten groups. We take one group as the test data set and use the remaining groups to train a model. Then, the candidate model is evaluated using the test data. In this way, we can get a predicted class label for all the samples. Then, the output of each classification experiment is compared to the ground truth, and the accuracy is computed to indicate how many class labels are correctly identified. The key parameters with the highest classification accuracies are selected.

Once we get an optimized PASCS-MP model, we can compare the performances of MMS, volume, and surface area measurements for classifying individuals of different Aβ status. We use the volume from the left and right hippocampi (i.e., hippocampi in each brain hemisphere) as two features to train the classifier instead of adding them together. The same classification strategy is applied to surface areas from both sides. Moreover, we will compare the classification performances based on PASCS-MP, PASS-MP (Zhang et al., 2017b, 2016b) and SPHARM (Chung et al., 2008, 2007; Shi et al., 2013a). We evaluate these classification performances with the same 10-fold cross-validation method. Four performance measures: the Accuracy (ACC), Balanced Accuracy (B-ACC), Specificity (SPE) and Sensitivity (SEN) are computed (Bhagwat et al., 2018; Hinrichs et al., 2011; Ritter et al., 2015; Salvatore et al., 2018; Zhang et al., 2017b). We also compute the area-under-the-curve (AUC) of the receiver operating characteristic (ROC) (Bhagwat et al., 2018; Fan et al., 2008; La Joie et al., 2013; Nakamura et al., 2018). By considering these performance measures, we expect the proposed system integrating MMS, PASCS-MP and the binary random forest classifier to perform better than similar classification strategies for identifying individuals with different Aβ status.

## 3. RESULTS

### 3.1 Key Parameter Estimations for the PASCS-MP Method

To apply PASCS-MP method on hippocampal MMS, four parameters need to be empirically assigned, namely: the patch size, the dimensionality of the learned sparse coding, the regularization parameter for the *l*_1_-norm (λ) and the kernel size (σ) in the exponential function. Selecting suitable parameters will lead to superior performance in refining lower dimensional MMS representations related to AD pathology. With 10-fold cross-validation, these key parameters are evaluated from PASCS-MP based classification performance on 109 AD patients and 180 CU subjects of ADNI-2 cohort. To avoid data leakage, these subjects are not used in the following study of Aβ burden classification.

In **Fig. 3**, we illustrate the classification accuracy for different values of each parameter. When we evaluate one parameter, we fix the rest parameters. For example, in the first bar chart in **Fig. 3**, we try different patch sizes including 5×5,10×10,15×15,20×20 and 30×30 while we fix the sparse code dimensionality as to 1800, and set λ to 0.22, and σ to 3.6. By testing varied sets of parameters, we find that the optimal patch size is 10×10, the optimal sparse code dimensionality is 1800, the optimal λ is 0.22 and the optimal σ is 3.6 and these optimal parameters will be adopted in the study of Aβ burden classification.

**Fig. 3.**
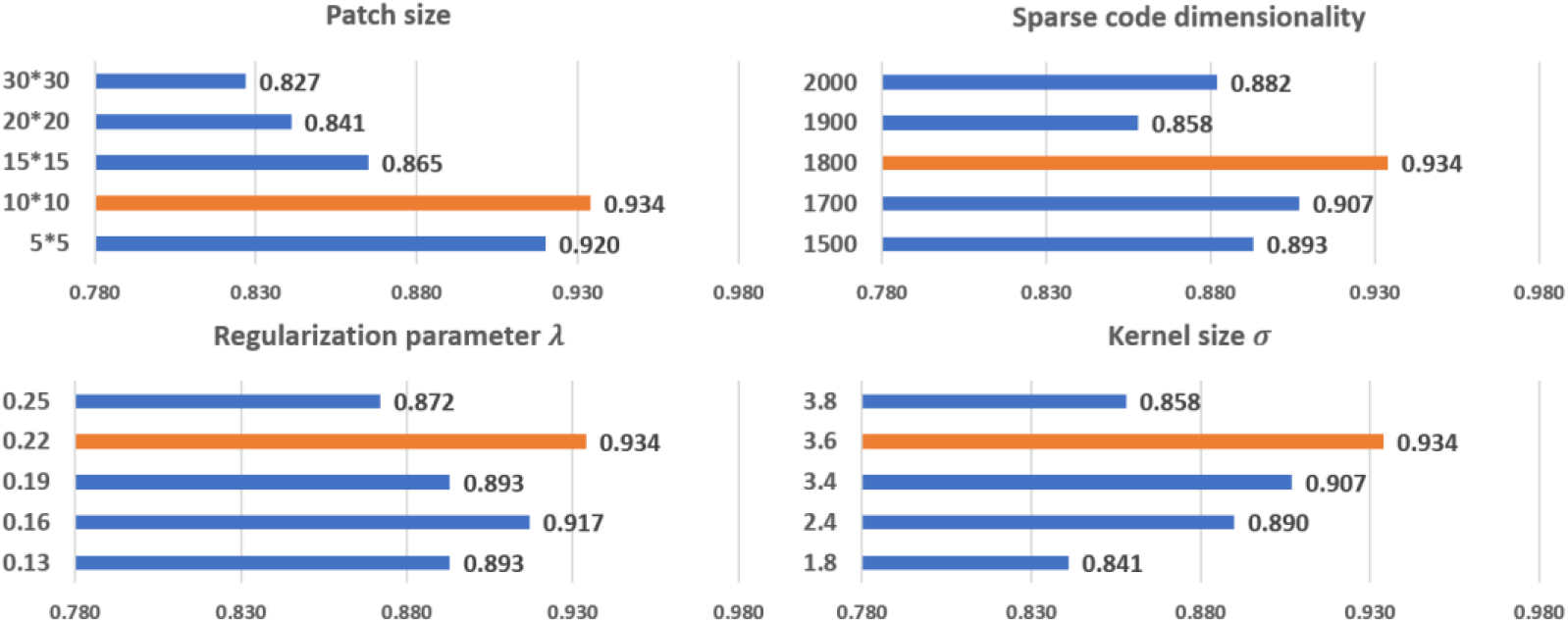
The relationship of each parameter to classification accuracy. The *y*-axis represents the value for each parameter. The orange bars represent the classification performances using the optimal parameters.

### 3.2 Classification of Aβ Burden

To explore whether there is a significant gain in classification power with our new system, based on our surface MMS, we generate two different kinds of sparse codes with our previous framework (PASS-MP) (Fu et al., 2021; Zhang et al., 2017; Zhang et al., 2016b) and the new framework (PASCS-MP). The parameter settings for the two sparse coding methods are the same. Additionally, we apply the popular SPHARM method (Chung et al., 2008; Shi et al., 2013a) to calculate hippocampal shape features. Based on these three kinds of feature sets, we apply the random forest classifier to detect individuals with different Aβ status. Moreover, we also examine the classification performances using hippocampal MMS, surface area and volume measures. These classification performances are evaluated using ACC, B-ACC, SPE, SEN. For each binary classification of ADNI cohort, we repeat the 10-fold cross-validation 5 times; the mean and 95% confident interval of the evaluation measures are calculated as (Vanwinckelen and Blockeel, 2012) and shown in the middle three columns of **Table 2**. To further evaluate the performance of our new framework, we firstly generate new representations with our proposed PASCS-MP for all the CU subjects from ADNI and OASIS cohorts. Then, we train a binary random forest model on the ADNI dataset and test it with the OASIS dataset. Since there is no cross-validation here, there is no confident interval in the last column of **Table 2**. We also compute the area-under-the-curve (AUC) of the receiver operating characteristic (ROC). The ROC curve and AUC for these classification tasks are illustrated in **Fig. 4**. This comparison analysis classification performance shows that the combination of PASCS-MP and hippocampal MMS measures have superior performance when detecting individuals with different Aβ status, compared to other similar methods.

**Table 2.** Classification Results for four contrasts.

| Area | Aβ+ AD vs.<br>Aβ- CU | Aβ+ MCI vs.<br>Aβ- MCI | Aβ+ CU vs. Aβ- CU<br>(ADNI) | Aβ+ CU vs. Aβ- CU<br>(OASIS) |
| --- | --- | --- | --- | --- |
| ACC | 0.68±0.01 | 0.55±0.02 | 0.54±0.01 | 0.47 |
| B-ACC | 0.69±0.02 | 0.55±0.02 | 0.54±0.02 | 0.43 |
| SPE | 0.66±0.02 | 0.54±0.02 | 0.55±0.02 | 0.49 |
| SEN | 0.71±0.03 | 0.56±0.03 | 0.53±0.04 | 0.37 |
| Volume | Aβ+ AD vs.<br>Aβ- CU | Aβ+ MCI vs.<br>Aβ- MCI | Aβ+ CU vs. Aβ- CU<br>(ADNI) | Aβ+ CU vs. Aβ- CU<br>(OASIS) |
| ACC | 0.71±0.01 | 0.53±0.02 | 0.50±0.03 | 0.51 |
| B-ACC | 0.72±0.01 | 0.53±0.01 | 0.50±0.03 | 0.52 |
| SPE | 0.68±0.01 | 0.52±0.01 | 0.51±0.02 | 0.54 |
| SEN | 0.75±0.01 | 0.54±0.02 | 0.49±0.04 | 0.50 |
| SPHARM | Aβ+ AD vs.<br>Aβ- CU | Aβ+ MCI vs.<br>Aβ- MCI | Aβ+ CU vs. Aβ- CU<br>(ADNI) | Aβ+ CU vs. Aβ- CU<br>(OASIS) |
| ACC | 0.71±0.02 | 0.56±0.02 | 0.52±0.02 | 0.60 |
| B-ACC | 0.71±0.02 | 0.56±0.03 | 0.51±0.04 | 0.60 |
| SPE | 0.74±0.02 | 0.61±0.03 | 0.56±0.03 | 0.61 |
| SEN | 0.68±0.04 | 0.51±0.03 | 0.46±0.05 | 0.60 |
| PASS-MP | Aβ+ AD vs.<br>Aβ- CU | Aβ+ MCI vs.<br>Aβ- MCI | Aβ+ CU vs. Aβ- CU<br>(ADNI) | Aβ+ CU vs. Aβ- CU<br>(OASIS) |
| ACC | 0.79±0.01 | 0.73±0.02 | 0.71±0.02 | 0.74 |
| B-ACC | 0.79±0.01 | 0.73±0.02 | 0.70±0.03 | 0.73 |
| SPE | 0.78±0.02 | 0.75±0.02 | 0.73±0.03 | 0.74 |
| SEN | 0.79±0.01 | 0.72±0.03 | 0.67±0.03 | 0.73 |
| PASCS-MP | Aβ+ AD vs.<br>Aβ- CU | Aβ+ MCI vs.<br>Aβ- MCI | Aβ+ CU vs. Aβ- CU<br>(ADNI) | Aβ+ CU vs. Aβ- CU<br>(OASIS) |
| ACC | **0.91±0.01** | **0.89±0.01** | **0.79±0.02** | **0.81** |
| B-ACC | **0.91±0.01** | **0.89±0.01** | **0.79±0.03** | **0.80** |
| SPE | **0.91±0.01** | **0.91±0.01** | **0.80±0.02** | **0.82** |
| SEN | **0.90±0.01** | **0.88±0.01** | **0.79±0.05** | **0.79** |

Values are mean ± 95% confident interval where applicable.

**Fig. 4.**
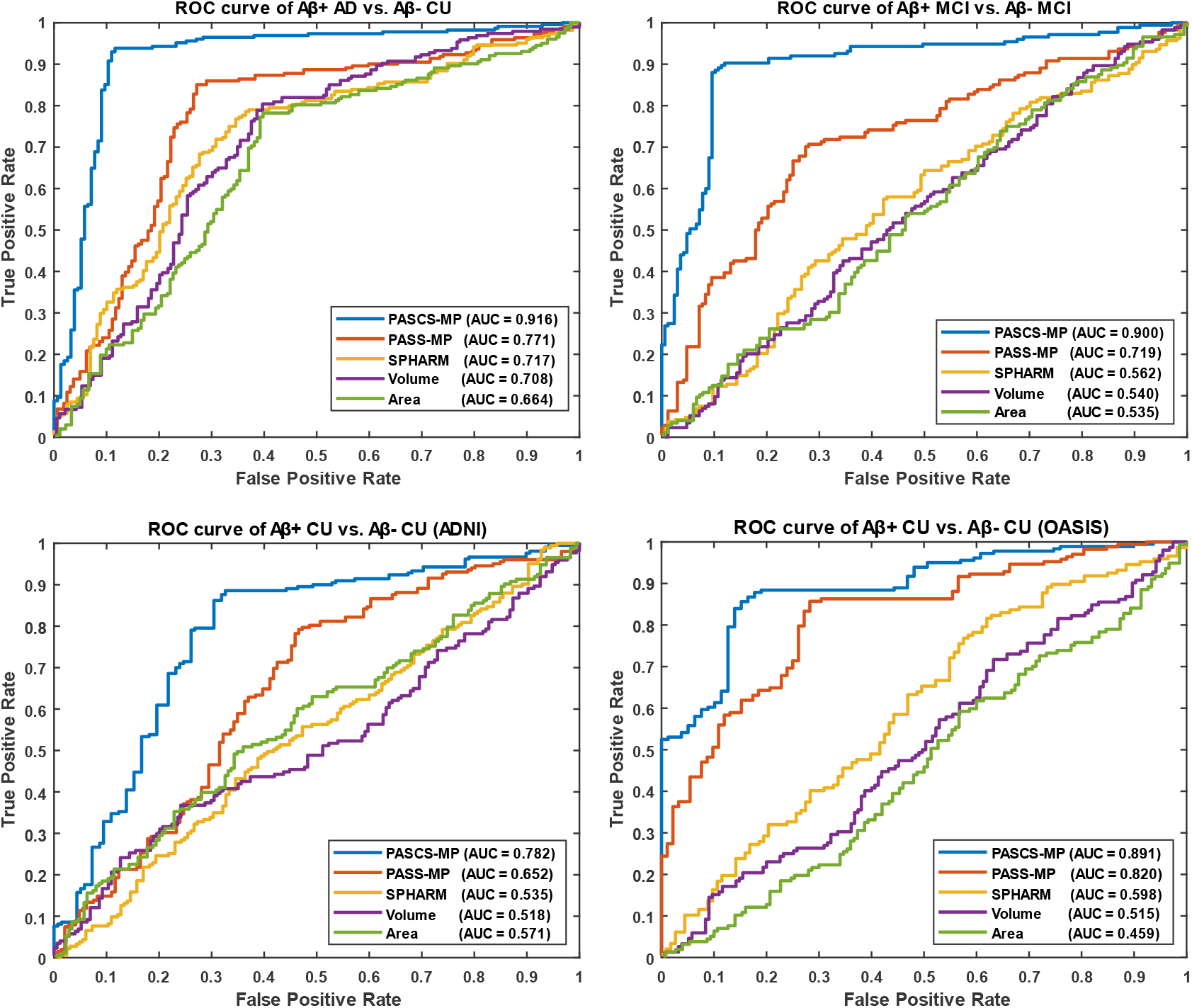
ROC curves for the classification tasks, Aβ+ AD vs. Aβ-CU, Aβ+ MCI vs. Aβ-MCI, Aβ+ CU vs. Aβ-CU (ADNI), and Aβ+ CU vs. Aβ-CU (OASIS).

## 4. DISCUSSION

In this paper, we propose a novel surface feature dimension reduction scheme, PASCS-MP, to correlate the hippocampus MMS with different levels of Aβ burden in individual subjects. We develop a hippocampal structure-based Aβ burden prediction system that involves hippocampal MMS computing, sparse coding and classification modules. We apply the proposed system on two independent datasets, ADNI and OASIS. We have two main findings. Firstly, the hippocampal surface-based MMS measure practically encodes a great deal of neighboring intrinsic geometry information that would otherwise be inaccessible or overlooked in classical hippocampal volume and surface area measures. Experimental results show that the MMS measure provides better classification accuracy than hippocampal volume, surface area measures and SPHARM for detecting the relationships between hippocampal deformations and Aβ positivity. Secondly, we propose a novel sparse coding method, PASCS-MP. It has all the advantages of our previous proposed PASS-MP (Zhang et al., 2016b, 2016a) and improves the follow-up classification performance compared to PASS-MP.

### 4.1 Comparison Analysis of Hippocampal MMS, Volume and Surface Area

The hippocampus is a primary target region for studying early AD progression. Its structure can be measured using the widely used overall hippocampal volume, surface area and our proposed hippocampal MMS. Our prior studies (Dong et al., 2019; Li et al., 2016; Shi et al., 2011; Wang et al., 2011) show that hippocampal MMS performs robustly in distinguishing clinical groups at different AD risk levels. In particular, we previously found that hippocampal MMS can detect *APOE4* gene dose effects on the hippocampus during the preclinical stage, while the hippocampal volume measure cannot (Dong et al., 2019). A study by Wu et al. (2018) demonstrates that hippocampal MMS performs better than traditional hippocampal volume measures in classifying 151 Aβ+ AD and 271 Aβ-CU subjects.

This work evaluates the performance of the above three hippocampal measurements for predicting Aβ status at the individual subject level. Classification results (see **Table 2** and **Fig. 4**) show that hippocampal MMS has better performance as measured by ACC, SPE, SEN and AUC. These results validate our hypothesis that hippocampal MMS-based analysis methods provide improved statistical accuracy than hippocampal volume and surface area measures in predicting the subjects with different Aβ status in the AD continuum. Our prior work (Wang et al., 2011) shows that MMS may offer a surrogate biomarker for PET/CSF Aβ biomarkers. This work further shows it can be used to classify brain Aβ burden on an individual basis.

### 4.2 Comparative Analysis of PASCS-MP, PASS-MP and SPHARM

The MMS measure for brain structures performs well in clinical group comparisons (Dong et al., 2020b, 2019; Li et al., 2016; Shi et al., 2015, 2014b; Wang et al., 2013; Yao et al., 2018), and as we have shown, it has the potential to further be applied for individual Aβ classification. To achieve this goal, we need to solve the challenge that the MMS dimension is usually much larger than the number of subjects, i.e., the so-called *high dimension, small sample size problem*. A reasonable solution is to reduce the feature dimension. Existing feature dimension reduction approaches include feature selection (Fan et al., 2005; Jain and Zongker, 1997), feature extraction (Guyon et al., 2008; Jolliffe, 2002; Mika et al., 1999) and sparse learning methods (Donoho, 2006; Vounou et al., 2010; Wang et al., 2013). In most cases, information is lost when mapping data into a lower-dimensional space. By defining a better lower-dimensional subspace, this information loss can be limited. Sparse coding (Lee et al., 2007; Mairal et al., 2009) has been previously proposed to learn an over-complete set of basis vectors (also called a *dictionary*) to represent input vectors efficiently and concisely (Donoho and Elad, 2003). Sparse coding has been shown to be effective for many tasks such as image imprinting (Moody et al., 2012), image deblurring (Yin et al., 2008), super-resolution (Yang et al., 2008), classification (Mairal et al., 2009), functional brain connectivity (Lv et al., 2017, 2015), and structural morphometry analysis (Zhang et al., 2017).

Our previous studies (Zhang et al., 2017; Zhang et al., 2016b, 2016a) propose a patch analysis-based surface sparse-coding and max-pooling (PASS-MP) method, consisting of sparse coding (Lee et al., 2006; Mairal et al., 2009) and max-pooling (LeCun et al., 2015), for surface feature dimension reduction. PASS-MP has excellent impressive performance for the sparse coding of our MMS features. Our prior studies successfully apply these sparse codes in detecting individual brain structure abnormalities and obtain state-of-art performance (Dong et al., 2020a; Fu et al., 2021; Wu et al., 2018).

Even so, there typically exists non-Gaussian and localized sources of noise in surface-based morphometry features, this can dramatically influence the learned dictionary and further lead to poor sparse coding based on the loss function of PASS-MP. The correntropy measure is a very robust method for correcting such sources of noise (He et al., 2012; Liu et al., 2007; Nikolova and Ng, 2006). In this paper, we improve upon the PASS-MP method by introducing correntropy measures into the loss function (Gui et al., 2017). Therefore, our proposed sparse coding method, PASCS-MP, incorporates all the advantages of PASS-MP and meanwhile improves the classification performance. We also test SPHARM-based hippocampal shape features as they have frequently been studied in prior AD research (e.g., (Cuingnet et al., 2011; Gerardin et al., 2009; Gutman et al., 2013)). In such an approach, we use a series of spherical harmonics to model the shapes of the hippocampus segmented by FSL. The SPHARM coefficients are computed using SPHARM-PDM (Spherical Harmonics-Point Distribution Model) software developed by the University of North Carolina and the National Alliance for Medical Imaging Computing (Styner et al., 2006). The classification features are based on these SPHARM coefficients, which are represented by two sets of three-dimensional SPHARM coefficients for each subject (in fact, one set for the hippocampus in each brain hemisphere). In Gerardin et al. (2009), they use a feature selection step because the subject groups are much smaller (fewer than 30 subjects in each group). When the number of subjects is small, the classifier can be more sensitive to uninformative features. In the current study, the number of subjects is relatively large, so a feature selection step is less necessary and may increase the risk of overfitting. We adopt the same approach in Cuingnet et al. (2011), who chose to avoid this selection step. The classification results (see **Table 2** and **Fig. 4**) based on PASCS-MP, PASS-MP and SPHARM meet our expectation that the classification performances based on PASCS-MP have an apparent improvement measured by ACC, B-ACC, SPE, SEN and AUC.

### 4.3 Aβ Burden Prediction using MRI Biomarkers

Aβ accumulation is a major feature of AD neuropathology (Brier et al., 2016; Cummings, 2019). Detecting it early and accurately provides a potential opportunity for effective therapeutic interventions before the advanced stages of AD (Tosun et al., 2014). Compared to PET and CSF Aβ measurement techniques, MRI is less expensive (than PET) and less invasive (than both PET and lumbar puncture). AD-related biomarker studies (Jack et al., 2018; Jack and Holtzman, 2013; Sperling et al., 2011b) have shown that abnormal brain Aβ accumulation typically precedes detectable structural brain abnormalities. There is emerging literature using MRI biomarkers to predict brain Aβ burden, and hippocampal structural measurement is one of the major predictors (Ansart et al., 2020; Pekkala et al., 2020; Tosun et al., 2016, 2014). Tosun et al. (2014) applied LASSO penalized logistic regression classifier to MRI-based voxel-wise anatomical shape variation measures and cerebral blood flow measures to predict Aβ positivity in 67 people with early MCI (34 Aβ+); the classification accuracy was 83%. Ansart et al. (2020) applied LASSO feature selection and a random forest classifier to MRI-based cortical thickness and hippocampal volume measures to classify 596 people with MCI scanned as part of ADNI MCI (375 Aβ+); the AUC was 0.80. Our proposed classification framework has a higher ACC=89% or AUC=0.90 than each of these two studies (Ansart et al., 2020; Tosun et al., 2014) for predicting Aβ status in people with MCI. Of the studies predicting Aβ positivity in CUs, Ansart et al. (2020) applied LASSO feature selection and random forest classifier to MRI-derived cortical thickness and hippocampal volume measures to classify 431 ADNI CUs (162 Aβ+) and 318 INSIGHT CUs (88 Aβ+); the AUCs were 0.59 and 0.62, respectively. Pekkala et al. (2020) used the Disease State Index machine learning algorithm and MRI-based biomarkers (total cortical and gray matter volumes, hippocampus, accumbens, thalamus and putament volumes) to predict Aβ burden in 48 CUs (20 Aβ+); the AUC was 0.78. Our proposed classification framework has AUC=0.78 on 348 ADNI CUs (116 Aβ+) and AUC=0.89 on 260 OASIS CUs (52 Aβ+). **Table 3** and **Fig. 4** present the AUC or ACC values from this work and from similar studies predicting Aβ positivity using brain MRI biomarkers. Compared to these similar studies, our proposed classification system only uses hippocampal structural features but still consistently outperforms other recently published methods for predicting Aβ positivity in people with MCI and CUs.

**Table 3.** Studies to impute Aβ status from MRI biomarkers in key clinical groups in AD research.

| Method | Subjects (Aβ+/-) | MRI Biomarkers | ACC | AUC |
| --- | --- | --- | --- | --- |
| PASCS-MP-Random forest classifier<br>(This work) | 342 ADNI MCI (171/171)<br>348 ADNI CU (116/232)<br>260 OASIS CU (52/208) | Hippocampal multivariate morphometry statistics (MMS) | 0.89±0.01<br>0.79±0.02<br>0.81 | 0.90<br>0.78<br>0.89 |
| LASSO penalized logistic regression classifier<br>(Tosun et al., 2014) | 67 Early MCI (34/33) | Voxel-wise anatomical shape variation measures and cerebral blood flow (including frontoparietal cortical, hippocampal regions, among others) | 0.83±0.03 | |
| LASSO feature selection and Random Forest classifier<br>(Ansart et al., 2020) | 596 ADNI MCI (375/221)<br>431 ADNI CU (162/269)<br>318 INSIGHT CU (88/230) | Cortical thickness and hippocampal volume |  | 0.80<br>0.59<br>0.62 |
| Disease State Index machine learning algorithm<br>(Pekkala et al., 2020) | 48 CU (20/28) | Total cortical and gray matter volumes, hippocampus, accumbens, thalamus and putamen volumes |  | 0.78 |

### 4.4 Limitations and Future Work

Despite the promising results are obtained by applying our proposed Aβ positivity classification framework, there are two important caveats. First, when applying the PASCS-MP method to refine MMS, we generally cannot visualize the selected features. To some extent, this decreases the interpretability of the effects, although it is still possible to visualize statistically significant regions as in our prior group difference studies (Shi et al., 2013b; Wang et al., 2013). However, in our recent work (Zhang et al., 2018), instead of randomly selecting patches to build the initial dictionary, we use group lasso screening to select the most significant features first. Therefore, the features used in sparse coding may be visualized on the surface map. In the future, we will incorporate this idea into the PASCS-MP framework to make it more interpretable. Second, this work only applies hippocampal MMS to predict Aβ positivity. In future work, we plan to introduce more AD risk factors (such as demographic information, genetic information and clinical assessments) (Ansart et al., 2020; Pekkala et al., 2020; Tosun et al., 2014), and more AD regions of interest (ROIs, e.g., ventricles, entorhinal cortex, temporal lobes) (Brier et al., 2016; Dong et al., 2020b; Foley et al., 2017) into our proposed framework; these additional features are expected to improve the Aβ positivity prediction.

## 5. CONCLUSION

In this paper, we explore the association between hippocampal structures and Aβ positivity on two independent databases using our hippocampal MMS, PASCS-MP method and a random forest classifier. Compared to traditional hippocampal shape measures, MMS have superior performance for predicting Aβ positivity in the AD continuum. Besides, the proposed PASCS-MP outperforms our previous sparse coding method (PASS-MP) on refining MMS features. Compared to similar studies, this work achieves state-of-the-art performance when predicting Aβ positivity based on MRI biomarkers. In future, we plan to apply this proposed framework to other AD ROIs and further improve the comprehensibility of the framework by visualizing morphometry features selected in the classification.

## ACKNOWLEDGMENTS

Algorithm development and image analysis for this study were partially supported by the National Institute on Aging (RF1AG051710, R21AG065942, R01AG031581 and P30AG19610), the National Library of Medicine, National Cancer Institute, the National Institute for Biomedical Imaging and Bioengineering (R01EB025032), and the Arizona Alzheimer Consortium.

Data collection and sharing for this project was funded by the Alzheimer’s Disease Neuroimaging Initiative (ADNI) (National Institutes of Health Grant U01 AG024904) and DoD ADNI (Department of Defense award number W81XWH-12-2-0012). ADNI is funded by the National Institute on Aging, the National Institute of Biomedical Imaging and Bioengineering, and through generous contributions from the following: Alzheimer’s Association; Alzheimer’s Drug Discovery Foundation; BioClinica, Inc.; Biogen Idec Inc.; Bristol-Myers Squibb Company; Eisai Inc.; Elan Pharmaceuticals, Inc.; Eli Lilly and Company; F. Hoffmann-La Roche Ltd and its affiliated company Genentech, Inc.; GE Healthcare; Innogenetics, N.V.; IXICO Ltd.; Janssen Alzheimer Immunotherapy Research & Development, LLC.; Johnson & Johnson Pharmaceutical Research & Development LLC.; Medpace, Inc.; Merck & Co., Inc.; Meso Scale Diagnostics, LLC.; NeuroRx Research; Novartis Pharmaceuticals Corporation; Pfizer Inc.; Piramal Imaging; Servier; Synarc Inc.; and Takeda Pharmaceutical Company. The Canadian Institutes of Health Research is providing funds to support ADNI clinical sites in Canada. Private sector contributions are facilitated by the Foundation for the National Institutes of Health (www.fnih.org). The grantee organization is the Northern California Institute for Research and Education, and the study is coordinated by the Alzheimer’s Disease Cooperative Study at the University of California, San Diego. ADNI data are disseminated by the Laboratory for Neuro Imaging at the University of Southern California.

## Footnotes

1 http://gsl.lab.asu.edu/software/pass-mp/

## Notes

### Competing Interest Statement

The authors have declared no competing interest.

### Summary of Updates

We update the experiment about the OASIS dataset. We train the model with ADNI and test it with OASIS.

## REFERENCES

Ansart, M., Epelbaum, S., Gagliardi, G., Colliot, O., Dormont, D., Dubois, B., Hampel, H., Durrleman, S., 2020. Reduction of recruitment costs in preclinical AD trials: validation of automatic pre-screening algorithm for brain amyloidosis. Stat. Methods Med. Res. 29, 151–164. https://doi.org/10.1177/0962280218823036

Apostolova, L.G., Morra, J.H., Green, A.E., Hwang, K.S., Avedissian, C., Woo, E., Cummings, J.L., Toga, A.W., Jack Jr., C.R., Weiner, M.W., Thompson, P.M., 2010. Automated 3D mapping of baseline and 12-month associations between three verbal memory measures and hippocampal atrophy in 490 ADNI subjects. Neuroimage 51, 488–499. https://doi.org/10.1016/j.neuroimage.2009.12.125

Bateman, R.J., Blennow, K., Doody, R., Hendrix, S., Lovestone, S., Salloway, S., Schindler, R., Weiner, M., Zetterberg, H., Aisen, P., Vellas, B., 2019. Plasma Biomarkers of AD Emerging as Essential Tools for Drug Development: An EU/US CTAD Task Force Report. J. Prev. Alzheimer’s Dis. 6, 169–173. https://doi.org/10.14283/jpad.2019.21

Bhagwat, N., Viviano, J.D., Voineskos, A.N., Chakravarty, M.M., 2018. Modeling and prediction of clinical symptom trajectories in Alzheimer’s disease using longitudinal data. PLoS Comput. Biol. 14, 1–25. https://doi.org/10.1371/journal.pcbi.1006376

Boureau, Y.L., Ponce, J., Lecun, Y., 2010. A theoretical analysis of feature pooling in visual recognition, in: ICML 2010 - Proceedings, 27th International Conference on Machine Learning.

Brier, M.R., Gordon, B., Friedrichsen, K., McCarthy, J., Stern, A., Christensen, J., Owen, C., Aldea, P., Su, Y., Hassenstab, J., Cairns, N.J., Holtzman, D.M., Fagan, A.M., Morris, J.C., Benzinger, T.L.S., Ances, B.M., 2016. Tau and Ab imaging, CSF measures, and cognition in Alzheimer’s disease. Sci. Transl. Med. 8, 1–10. https://doi.org/10.1126/scitranslmed.aaf2362

Bro-Nielsen, M., Gramkow, C., 1996. Fast fluid registration of medical images, in: Lecture Notes in Computer Science (Including Subseries Lecture Notes in Artificial Intelligence and Lecture Notes in Bioinformatics). https://doi.org/10.1007/bfb0046964

Brookmeyer, R., Johnson, E., Ziegler-Graham, K., Arrighi, H.M., 2007. Forecasting the global burden of Alzheimer’s disease. Alzheimer’s Dement. 3, 186–191. https://doi.org/10.1016/j.jalz.2007.04.381

Cacciaglia, R., Molinuevo, J.L., Falcón, C., Brugulat-Serrat, A., Sánchez-Benavides, G., Gramunt, N., Esteller, M., Morán, S., Minguillón, C., Fauria, K., Gispert, J.D., 2018. Effects of APOE -∊4 allele load on brain morphology in a cohort of middle-aged healthy individuals with enriched genetic risk for Alzheimer’s disease. Alzheimer’s Dement. 14, 902–912. https://doi.org/10.1016/j.jalz.2018.01.016

Ching, C.R.K., Abaryan, Z., Santhalingam, V., Zhu, A.H., Bright, J.K., Jahanshad, N., Thompson, P.M., 2020. Sex-Dependent Age Trajectories of Subcortical Brain Structures: Analysis of Large-Scale Percentile Models and Shape Morphometry. bioRxiv 2020.09.30.321711. https://doi.org/10.1101/2020.09.30.321711

Chung, M.K., Dalton, K.M., Davidson, R.J., 2008. Tensor-based cortical surface morphometry via weighted spherical harmonic representation. IEEE Trans. Med. Imaging. https://doi.org/10.1109/TMI.2008.918338

Chung, M.K., Dalton, K.M., Shen, L., Evans, A.C., Davidson, R.J., 2007. Weighted Fourier Series Representation and Its Application to Quantifying the Amount of Gray Matter. IEEE Trans. Med. Imaging 26, 566–581. https://doi.org/10.1109/TMI.2007.892519

Colom, R., Stein, J.L., Rajagopalan, P., Martínez, K., Hermel, D., Wang, Y., Álvarez-Linera, J., Burgaleta, M., Quiroga, M.Á., Shih, P.C., Thompson, P.M., 2013. Hippocampal structure and human cognition: Key role of spatial processing and evidence supporting the efficiency hypothesis in females. Intelligence. https://doi.org/10.1016/j.intell.2013.01.002

Costafreda, S.G., Dinov, I.D., Tu, Z., Shi, Y., Liu, C.-Y., Kloszewska, I., Mecocci, P., Soininen, H., Tsolaki, M., Vellas, B., others, 2011. Automated hippocampal shape analysis predicts the onset of dementia in mild cognitive impairment. Neuroimage 56, 212–219.

Crivello, F., Lemaître, H., Dufouil, C., Grassiot, B., Delcroix, N., Tzourio-Mazoyer, N., Tzourio, C., Mazoyer, B., 2010. Effects of ApoE-ɛ4 allele load and age on the rates of grey matter and hippocampal volumes loss in a longitudinal cohort of 1186 healthy elderly persons. Neuroimage 53, 1064–1069. https://doi.org/10.1016/j.neuroimage.2009.12.116

Cuingnet, R., Gerardin, E., Tessieras, J., Auzias, G., Lehéricy, S., Habert, M.O., Chupin, M., Benali, H., Colliot, O., 2011. Automatic classification of patients with Alzheimer’s disease from structural MRI: A comparison of ten methods using the ADNI database. Neuroimage 56, 766–781. https://doi.org/10.1016/j.neuroimage.2010.06.013

Cummings, J., 2019. The National Institute on Aging—Alzheimer’s Association Framework on Alzheimer’s disease: Application to clinical trials. Alzheimer’s Dement. 15, 172–178. https://doi.org/10.1016/j.jalz.2018.05.006

D’Agostino, E., Maes, F., Vandermeulen, D., Suetens, P., 2003. A viscous fluid model for multimodal non-rigid image registration using mutual information. Med. Image Anal. https://doi.org/10.1016/S1361-8415(03)00039-2

Davatzikos, C., Resnick, S.M., Wu, X., Parmpi, P., Clark, C.M., 2008. Individual patient diagnosis of AD and FTD via high-dimensional pattern classification of MRI. Neuroimage. https://doi.org/10.1016/j.neuroimage.2008.03.050

Dong, Q., Zhang, J., Li, Q., Wang, J., Leporé, N., Thompson, P.M., Caselli, R.J., Ye, J., Wang, Y., 2020a. Integrating Convolutional Neural Networks and Multi-Task Dictionary Learning for Cognitive Decline Prediction with Longitudinal Images. J. Alzheimers. Dis. https://doi.org/10.3233/JAD-190973

Dong, Q., Zhang, W., Stonnington, C.M., Wu, J., Gutman, B.A., Chen, K., Su, Y., Baxter, L.C., Thompson, P.M., Reiman, E.M., Caselli, R.J., Wang, Y., 2020b. Applying surface-based morphometry to study ventricular abnormalities of cognitively unimpaired subjects prior to clinically significant memory decline. NeuroImage Clin. 27. https://doi.org/10.1016/j.nicl.2020.102338

Dong, Q., Zhang, W., Wu, J., Li, B., Schron, E.H., McMahon, T., Shi, J., Gutman, B.A., Chen, K., Baxter, L.C., Thompson, P.M., Reiman, E.M., Caselli, R.J., Wang, Y., 2019. Applying surface-based hippocampal morphometry to study APOE-E4 allele dose effects in cognitively unimpaired subjects. NeuroImage Clin. https://doi.org/10.1016/j.nicl.2019.101744

Donoho, D.L., 2006. Compressed sensing. IEEE Trans. Inf. Theory. https://doi.org/10.1109/TIT.2006.871582

Donoho, D.L., Elad, M., 2003. Optimally sparse representation in general (nonorthogonal) dictionaries via l1 minimization. Proc. Natl. Acad. Sci. 100, 2197–2202.

Dubey, R., Zhou, J., Wang, Y., Thompson, P.M., Ye, J., 2014. Analysis of sampling techniques for imbalanced data: An n=648 ADNI study. Neuroimage 87, 220–241. https://doi.org/10.1016/j.neuroimage.2013.10.005

Fan, Y., Resnick, S.M., Wu, X., Davatzikos, C., 2008. Structural and functional biomarkers of prodromal Alzheimer’s disease: A high-dimensional pattern classification study. Neuroimage 41, 277–285. https://doi.org/10.1016/j.neuroimage.2008.02.043

Fan, Y., Shen, D., Davatzikos, C., 2005. {C}lassification of structural images via high-dimensional image warping, robust feature extraction, and {S}{V}{M}. Med Image Comput Comput Assist Interv 8, 1–8.

Feng, Y., Huang, X., Shi, L., Yang, Y., Suykens, J.A.K., 2015. Learning with the maximum correntropy criterion induced losses for regression. J. Mach. Learn. Res. 16, 993–1034.

Fleisher, A.S., Chen, K., Liu, X., Roontiva, A., Thiyyagura, P., Ayutyanont, N., Joshi, A.D., Clark, C.M., Mintun, M.A., Pontecorvo, M.J., Doraiswamy, P.M., Johnson, K.A., Skovronsky, D.M., Reiman, E.M., 2011. Using positron emission tomography and florbetapir F 18 to image cortical amyloid in patients with mild cognitive impairment or dementia due to Alzheimer disease. Arch. Neurol. https://doi.org/10.1001/archneurol.2011.150

Foley, S.F., Tansey, K.E., Caseras, X., Lancaster, T., Bracht, T., Parker, G., Hall, J., Williams, J., Linden, D.E., 2017. Multimodal Brain Imaging Reveals Structural Differences in Alzheimer’s Disease Polygenic Risk Carriers: A Study in Healthy Young Adults. Biol Psychiatry 81, 154–161. https://doi.org/10.1016/j.biopsych.2016.02.033

Folstein, M.F., Folstein, S.E., McHugh, P.R., 1975. “Mini-mental state”. A practical method for grading the cognitive state of patients for the clinician. J. Psychiatr. Res. https://doi.org/10.1016/0022-3956(75)90026-6

Fu, W.J., 1998. Penalized regressions: The bridge versus the lasso? J. Comput. Graph. Stat. https://doi.org/10.1080/10618600.1998.10474784

Fu, Y., Zhang, J., Li, Yuan, Shi, J., Zou, Y., Guo, H., Li, Yongchao, Yao, Z., Wang, Y., Hu, B., 2021. A novel pipeline leveraging surface-based features of small subcortical structures to classify individuals with autism spectrum disorder. Prog. Neuro-Psychopharmacology Biol. Psychiatry 104, 109989. https://doi.org/10.1016/j.pnpbp.2020.109989

Gerardin, E., Chételat, G., Chupin, M., Cuingnet, R., Desgranges, B., Kim, H.-S., Niethammer, M., Dubois, B., Lehéricy, S., Garnero, L., Eustache, F., Colliot, O., 2009. Multidimensional classification of hippocampal shape features discriminates Alzheimer’s disease and mild cognitive impairment from normal aging. Neuroimage 47, 1476–1486. https://doi.org/10.1016/j.neuroimage.2009.05.036

Gui, J., Sun, Z., Ji, S., Tao, D., Tan, T., 2017. Feature selection based on structured sparsity: a comprehensive study. IEEE Trans. Neural Networks Learn. Syst. https://doi.org/10.1109/TNNLS.2016.2551724

Gutman, B.A., Hua, X., Rajagopalan, P., Chou, Y.Y., Wang, Y., Yanovsky, I., Toga, A.W., Jack, C.R., Weiner, M.W., Thompson, P.M., 2013. Maximizing power to track Alzheimer’s disease and MCI progression by LDA-based weighting of longitudinal ventricular surface features. Neuroimage 70, 386–401. https://doi.org/10.1016/j.neuroimage.2012.12.052

Guyon, I., Gunn, S., Nikravesh, M., Zadeh, L.A., 2008. Feature extraction: foundations and applications. Springer.

Han, X., Xu, C., Prince, J.L., 2003. A topology preserving level set method for geometric deformable models. IEEE Trans. Pattern Anal. Mach. Intell. https://doi.org/10.1109/TPAMI.2003.1201824

Hardy, J., Selkoe, D.J., 2002. The amyloid hypothesis of Alzheimer’s disease: progress and problems on the road to therapeutics. Science (80-.). 297, 353–356. https://doi.org/10.1126/science.1072994

He, R., Tan, T., Wang, L., Zheng, W.S., 2012. L 2, 1 regularized correntropy for robust feature selection, in: Proceedings of the IEEE Computer Society Conference on Computer Vision and Pattern Recognition. https://doi.org/10.1109/CVPR.2012.6247966

Hinrichs, C., Singh, V., Xu, G., Johnson, S.C., 2011. Predictive markers for AD in a multi-modality framework: An analysis of MCI progression in the ADNI population. Neuroimage 55, 574–589. https://doi.org/10.1016/j.neuroimage.2010.10.081

Hoppe, H., 1996. Progressive meshes, in: Proceedings of the 23rd Annual Conference on Computer Graphics and Interactive Techniques, SIGGRAPH 1996. https://doi.org/10.1145/237170.237216

Hyman, B.T., 2011. Amyloid-dependent and amyloid-independent stages of alzheimer disease. Arch. Neurol. https://doi.org/10.1001/archneurol.2011.70

Jack, C.R., Bennett, D.A., Blennow, K., Carrillo, M.C., Dunn, B., Haeberlein, S.B., Holtzman, D.M., Jagust, W., Jessen, F., Karlawish, J., Liu, E., Molinuevo, J.L., Montine, T., Phelps, C., Rankin, K.P., Rowe, C.C., Scheltens, P., Siemers, E., Snyder, H.M., Sperling, R., Elliott, C., Masliah, E., Ryan, L., Silverberg, N., 2018. NIA-AA Research Framework: Toward a biological definition of Alzheimer’s disease. Alzheimer’s Dement. 14, 535–562. https://doi.org/10.1016/j.jalz.2018.02.018

Jack, C.R., Bennett, D.A., Blennow, K., Carrillo, M.C., Feldman, H.H., Frisoni, G.B., Hampel, H., Jagust, W.J., Johnson, K.A., Knopman, D.S., Petersen, R.C., Scheltens, P., Sperling, R.A., Dubois, B., 2016. A/T/N: An unbiased descriptive classification scheme for Alzheimer disease biomarkers. Neurology 87, 539–547. https://doi.org/10.1212/WNL.0000000000002923

Jack, C.R., Holtzman, D.M., 2013. Biomarker modeling of alzheimer’s disease. Neuron 80, 1347–1358. https://doi.org/10.1016/j.neuron.2013.12.003

Jain, A., Zongker, D., 1997. Feature selection: Evaluation, application, and small sample performance. Pattern Anal. Mach. Intell. IEEE Trans. 19, 153–158.

Janelidze, S., Mattsson, N., Palmqvist, S., Smith, R., Beach, T.G., Serrano, G.E., Chai, X., Proctor, N.K., Eichenlaub, U., Zetterberg, H., Blennow, K., Reiman, E.M., Stomrud, E., Dage, J.L., Hansson, O., 2020. Plasma P-tau181 in Alzheimer’s disease: relationship to other biomarkers, differential diagnosis, neuropathology and longitudinal progression to Alzheimer’s dementia. Nat. Med. 26, 379–386. https://doi.org/10.1038/s41591-020-0755-1

Jianchao Yang, Wright, J., Huang, T., Yi Ma, 2008. Image super-resolution as sparse representation of raw image patches, in: 2008 IEEE Conference on Computer Vision and Pattern Recognition. IEEE, pp. 1–8. https://doi.org/10.1109/CVPR.2008.4587647

Jolliffe, I.T., 2002. Principal Component Analysis, Second Edition. Encycl. Stat. Behav. Sci. https://doi.org/10.2307/1270093

Kao, P.Y., Shailja, F., Jiang, J., Zhang, A., Khan, A., Chen, J.W., Manjunath, B.S., 2020. Improving Patch-Based Convolutional Neural Networks for MRI Brain Tumor Segmentation by Leveraging Location Information. Front. Neurosci. https://doi.org/10.3389/fnins.2019.01449

Klunk, W.E., Koeppe, R.A., Price, J.C., Benzinger, T.L., Devous, M.D., Jagust, W.J., Johnson, K.A., Mathis, C.A., Minhas, D., Pontecorvo, M.J., Rowe, C.C., Skovronsky, D.M., Mintun, M.A., 2015. The Centiloid project: Standardizing quantitative amyloid plaque estimation by PET. Alzheimer’s Dement. 11, 1–15.e4. https://doi.org/10.1016/j.jalz.2014.07.003

La Joie, R., Perrotin, A., De La Sayette, V., Egret, S., Doeuvre, L., Belliard, S., Eustache, F., Desgranges, B., Chételat, G., 2013. Hippocampal subfield volumetry in mild cognitive impairment, Alzheimer’s disease and semantic dementia. NeuroImage Clin. https://doi.org/10.1016/j.nicl.2013.08.007

Lee, H., Battle, A., Raina, R., Ng, A.Y., 2007. Efficient sparse coding algorithms, in: Advances in Neural Information Processing Systems. https://doi.org/10.7551/mitpress/7503.003.0105

Lee, Y.K., Hou, S.W., Lee, C.C., Hsu, C.Y., Huang, Y.S., Su, Y.C., 2013. Increased Risk of Dementia in Patients with Mild Traumatic Brain Injury: A Nationwide Cohort Study. PLoS One. https://doi.org/10.1371/journal.pone.0062422

Leow, A., Huang, S.C., Geng, A., Becker, J., Davis, S., Toga, A., Thompson, P., 2005. Inverse consistent mapping in 3D deformable image registration: Its construction and statistical properties, in: Lecture Notes in Computer Science. https://doi.org/10.1007/11505730_41

Li, B., Shi, J., Gutman, B.A., Baxter, L.C., Thompson, P.M., Caselli, R.J., Wang, Y., Neuroimaging Initiative, D., 2016. Influence of APOE Genotype on Hippocampal Atrophy over Time-An N=1925 Surface-Based ADNI Study. https://doi.org/10.1371/journal.pone.0152901

Liaw, A., Wiener, M., 2002. Classification and Regression by randomForest. R News.

Lin, B., Li, Q., Sun, Q., Lai, M.-J., Davidson, I., Fan, W., Ye, J., 2014. Stochastic Coordinate Coding and Its Application for Drosophila Gene Expression Pattern Annotation.

Liu, W., Pokharel, P.P., Principe, J.C., 2007. Correntropy: Properties and applications in non-Gaussian signal processing. IEEE Trans. Signal Process. https://doi.org/10.1109/TSP.2007.896065

Loop, C., 1987. Smooth Subdivision Surfaces Based on Triangles. Acm Siggraph.

Lorensen, W.E., Cline, H.E., 1987. Marching cubes: A high resolution 3D surface construction algorithm, in: Proceedings of the 14th Annual Conference on Computer Graphics and Interactive Techniques, SIGGRAPH 1987. https://doi.org/10.1145/37401.37422

Luders, E., Thompson, P.M., Kurth, F., Hong, J.Y., Phillips, O.R., Wang, Y., Gutman, B.A., Chou, Y.Y., Narr, K.L., Toga, A.W., 2013. Global and regional alterations of hippocampal anatomy in long-term meditation practitioners. Hum. Brain Mapp. https://doi.org/10.1002/hbm.22153

Lv, J., Jiang, X., Li, X., Zhu, D., Zhang, S., Zhao, S., Chen, H., Zhang, T., Hu, X., Han, J., Ye, J., Guo, L., Liu, T., 2015. {H}olistic atlases of functional networks and interactions reveal reciprocal organizational architecture of cortical function. IEEE Trans Biomed Eng 62, 1120–1131.

Lv, J., Lin, B., Li, Q., Zhang, W., Zhao, Y., Jiang, X., Guo, L., Han, J., Hu, X., Guo, C., Ye, J., Liu, T., 2017. Task fMRI data analysis based on supervised stochastic coordinate coding. Med. Image Anal. 38, 1–16. https://doi.org/10.1016/j.media.2016.12.003

Mairal, J., Bach, F., Ponce, J., Sapiro, G., 2009. Online dictionary learning for sparse coding, in: ACM International Conference Proceeding Series. https://doi.org/10.1145/1553374.1553463

Marcus, D.S., Fotenos, A.F., Csernansky, J.G., Morris, J.C., Buckner, R.L., 2010. Open Access Series of Imaging Studies: Longitudinal MRI Data in Nondemented and Demented Older Adults. J. Cogn. Neurosci. 22, 2677–2684. https://doi.org/10.1162/jocn.2009.21407

Mika, S., Ratsch, G., Weston, J., Scholkopf, B., Muller, K.R., 1999. Fisher discriminant analysis with kernels, in: Neural Networks for Signal Processing - Proceedings of the IEEE Workshop. https://doi.org/10.1109/nnsp.1999.788121

Monje, M., Thomason, M.E., Rigolo, L., Wang, Y., Waber, D.P., Sallan, S.E., Golby, A.J., 2013. Functional and structural differences in the hippocampus associated with memory deficits in adult survivors of acute lymphoblastic leukemia. Pediatr. Blood Cancer. https://doi.org/10.1002/pbc.24263

Moody, D.I., Brumby, S.P., Rowland, J.C., Gangodagamage, C., 2012. Unsupervised land cover classification in multispectral imagery with sparse representations on learned dictionaries, in: Proceedings - Applied Imagery Pattern Recognition Workshop. https://doi.org/10.1109/AIPR.2012.6528190

Morra, J.H., Tu, Z., Apostolova, L.G., Green, A.E., Avedissian, C., Madsen, S.K., Parikshak, N., Hua, X., Toga, A.W., Jack, C.R., Schuff, N., Weiner, M.W., Thompson, P.M., 2009. Automated 3D mapping of hippocampal atrophy and its clinical correlates in 400 subjects with Alzheimer’s disease, mild cognitive impairment, and elderly controls. Hum. Brain Mapp. 30, 2766–2788. https://doi.org/10.1002/hbm.20708

Mueller, S.G., Weiner, M.W., Thal, L.J., Petersen, R.C., Jack, C., Jagust, W., Trojanowski, J.Q., Toga, A.W., Beckett, L., 2005. The Alzheimer’s disease neuroimaging initiative. Neuroimaging Clin. N. Am. https://doi.org/10.1016/j.nic.2005.09.008

Nakamura, A., Kaneko, N., Villemagne, V.L., Kato, T., Doecke, J., Doré, V., Fowler, C., Li, Q.X., Martins, R., Rowe, C., Tomita, T., Matsuzaki, K., Ishii, Kenji, Ishii, Kazunari, Arahata, Y., Iwamoto, S., Ito, K., Tanaka, K., Masters, C.L., Yanagisawa, K., 2018. High performance plasma amyloid-β biomarkers for Alzheimer’s disease. Nature 554, 249–254. https://doi.org/10.1038/nature25456

Navitsky, M., Joshi, A.D., Kennedy, I., Klunk, W.E., Rowe, C.C., Wong, D.F., Pontecorvo, M.J., Mintun, M.A., Devous, M.D., 2018. Standardization of amyloid quantitation with florbetapir standardized uptake value ratios to the Centiloid scale. Alzheimer’s Dement. https://doi.org/10.1016/j.jalz.2018.06.1353

Nikolova, M., Ng, M.K., 2006. Analysis of half-quadratic minimization methods for signal and image recovery. SIAM J. Sci. Comput. https://doi.org/10.1137/030600862

Olshausen, B.A., Field, D.J., 1997. Sparse coding with an overcomplete basis set: A strategy employed by V1? Vision Res. https://doi.org/10.1016/S0042-6989(97)00169-7

Palmqvist, S., Janelidze, S., Quiroz, Y.T., Zetterberg, H., Lopera, F., Stomrud, E., Su, Y., Chen, Y., Serrano, G.E., Leuzy, A., Mattsson-Carlgren, N., Strandberg, O., Smith, R., Villegas, A., Sepulveda-Falla, D., Chai, X., Proctor, N.K., Beach, T.G., Blennow, K., Dage, J.L., Reiman, E.M., Hansson, O., 2020. Discriminative Accuracy of Plasma Phospho-tau217 for Alzheimer Disease vs Other Neurodegenerative Disorders. JAMA 324, 772. https://doi.org/10.1001/jama.2020.12134

Paquette, N., Shi, J., Wang, Y., Lao, Y., Ceschin, R., Nelson, M.D., Panigrahy, A., Lepore, N., 2017. Ventricular shape and relative position abnormalities in preterm neonates. NeuroImage. Clin. 15, 483–493. https://doi.org/10.1016/j.nicl.2017.05.025

Patenaude, B., Smith, S.M., Kennedy, D.N., Jenkinson, M., 2011. A Bayesian model of shape and appearance for subcortical brain segmentation. Neuroimage. https://doi.org/10.1016/j.neuroimage.2011.02.046

Pekkala, T., Hall, A., Ngandu, T., Gils, M. van, Helisalmi, S., Hänninen, T., Kemppainen, N., Liu, Y., Lötjönen, J., Paajanen, T., Rinne, J.O., Soininen, H., Kivipelto, M., Solomon, A., 2020. Detecting Amyloid Positivity in Elderly With Increased Risk of Cognitive Decline. Front. Aging Neurosci. 12, 1–9. https://doi.org/10.3389/fnagi.2020.00228

Pizer, S.M., Fritsch, D.S., Yushkevich, P.A., Johnson, V.E., Chaney, E.L., 1999. Segmentation, registration, and measurement of shape variation via image object shape. IEEE Trans. Med. Imaging. https://doi.org/10.1109/42.811263

Qiu, A., Taylor, W.D., Zhao, Z., MacFall, J.R., Miller, M.I., Key, C.R., Payne, M.E., Steffens, D.C., Krishnan, K.R., 2009. APOE related hippocampal shape alteration in geriatric depression. Neuroimage 44, 620–626. https://doi.org/S1053-8119(08)01121-X[pii]10.1016/j.neuroimage.2008.10.010 [doi]

Reiter, K., Nielson, K.A., Durgerian, S., Woodard, J.L., Smith, J.C., Seidenberg, M., Kelly, D.A., Rao, S.M., 2017. Five-Year Longitudinal Brain Volume Change in Healthy Elders at Genetic Risk for Alzheimer’s Disease. J. Alzheimer’s Dis. 55, 1363–1377. https://doi.org/10.3233/JAD-160504

Rey, D., Subsol, G., Delingette, H., Ayache, N., 2002. Automatic detection and segmentation of evolving processes in 3D medical images: Application to multiple sclerosis. Med. Image Anal. https://doi.org/10.1016/S1361-8415(02)00056-7

Ritter, K., Schumacher, J., Weygandt, M., Buchert, R., Allefeld, C., Haynes, J.D., 2015. Multimodal prediction of conversion to Alzheimer’s disease based onincomplete biomarkers. Alzheimer’s Dement. Diagnosis, Assess. Dis. Monit. 1, 206–215. https://doi.org/10.1016/j.dadm.2015.01.006

Rowe, C.C., Doré, V., Jones, G., Baxendale, D., Mulligan, R.S., Bullich, S., Stephens, A.W., De Santi, S., Masters, C.L., Dinkelborg, L., Villemagne, V.L., 2017. 18F-Florbetaben PET beta-amyloid binding expressed in Centiloids. Eur. J. Nucl. Med. Mol. Imaging 44, 2053–2059. https://doi.org/10.1007/s00259-017-3749-6

Salvatore, C., Cerasa, A., Castiglioni, I., 2018. MRI Characterizes the Progressive Course of AD and Predicts Conversion to Alzheimer’s Dementia 24 Months Before Probable Diagnosis. Front. Aging Neurosci. 10, 135. https://doi.org/10.3389/fnagi.2018.00135

Shen, L., Firpi, H.A., Saykin, A.J., West, J.D., 2009. Parametric surface modeling and registration for comparison of manual and automated segmentation of the hippocampus. Hippocampus 19, 588–595. https://doi.org/10.1002/hipo.20613

Shi, J., Leporé, N., Gutman, B.A., Thompson, P.M., Baxter, L.C., Caselli, R.J., Wang, Y., 2014. Genetic influence of apolipoprotein E4 genotype on hippocampal morphometry: An N = 725 surface-based Alzheimer’s disease neuroimaging initiative study. Hum. Brain Mapp. https://doi.org/10.1002/hbm.22447

Shi, J., Stonnington, C.M., Thompson, P.M., Chen, K., Gutman, B., Reschke, C., Baxter, L.C., Reiman, E.M., Caselli, R.J., Wang, Y., 2015. Studying ventricular abnormalities in mild cognitive impairment with hyperbolic Ricci flow and tensor-based morphometry. Neuroimage. https://doi.org/10.1016/j.neuroimage.2014.09.062

Shi, J., Thompson, P.M., Gutman, B., Wang, Y., 2013a. Surface fluid registration of conformal representation: Application to detect disease burden and genetic influence on hippocampus. Neuroimage. https://doi.org/10.1016/j.neuroimage.2013.04.018

Shi, J., Thompson, P.M., Wang, Y., 2011. Human Brain Mapping with Conformal Geometry and Multivariate Tensor-Based Morphometry, in: Lecture Notes in Computer Science (Including Subseries Lecture Notes in Artificial Intelligence and Lecture Notes in Bioinformatics). pp. 126–134. https://doi.org/10.1007/978-3-642-24446-9_16

Shi, J., Wang, Y., Ceschin, R., An, X., Lao, Y., Vanderbilt, D., Nelson, M.D., Thompson, P.M., Panigrahy, A., Leporé, N., 2013b. A Multivariate Surface-Based Analysis of the Putamen in Premature Newborns: Regional Differences within the Ventral Striatum. PLoS One. https://doi.org/10.1371/journal.pone.0066736

Sperling, Reisa A, Aisen, P.S., Beckett, L.A., Bennett, D.A., Craft, S., Fagan, A.M., Iwatsubo, T., Jack, C.R., Kaye, J., Montine, T.J., Park, D.C., Reiman, E.M., Rowe, C.C., Siemers, E., Stern, Y., Yaffe, K., Carrillo, M.C., Thies, B., Morrison-Bogorad, M., Wagster, M. V, Phelps, C.H., 2011. Toward defining the preclinical stages of Alzheimer’s disease: recommendations from the National Institute on Aging-Alzheimer’s Association workgroups on diagnostic guidelines for Alzheimer’s disease. Alzheimers. Dement. 7, 280–92. https://doi.org/10.1016/j.jalz.2011.03.003

Sperling, Reisa A., Jack, C.R., Black, S.E., Frosch, M.P., Greenberg, S.M., Hyman, B.T., Scheltens, P., Carrillo, M.C., Thies, W., Bednar, M.M., Black, R.S., Brashear, H.R., Grundman, M., Siemers, E.R., Feldman, H.H., Schindler, R.J., 2011. Amyloid-related imaging abnormalities in amyloid-modifying therapeutic trials: Recommendations from the Alzheimer’s Association Research Roundtable Workgroup. Alzheimer’s Dement. 7, 367–385. https://doi.org/10.1016/j.jalz.2011.05.2351

Styner, M., Lieberman, J.A., Pantazis, D., Gerig, G., 2004. Boundary and medial shape analysis of the hippocampus in schizophrenia. Med Image Anal 8, 197–203. https://doi.org/10.1016/j.media.2004.06.004

Styner, M., Oguz, I., Xu, S., Brechbühler, C., Pantazis, D., Levitt, J.J., Shenton, M.E., Gerig, G., 2006. Framework for the Statistical Shape Analysis of Brain Structures using SPHARM-PDM. Insight J. 242–250.

Su, Y., Blazey, T.M., Snyder, A.Z., Raichle, M.E., Marcus, D.S., Ances, B.M., Bateman, R.J., Cairns, N.J., Aldea, P., Cash, L., Christensen, J.J., Friedrichsen, K., Hornbeck, R.C., Farrar, A.M., Owen, C.J., Mayeux, R., Brickman, A.M., Klunk, W., Price, J.C., Thompson, P.M., Ghetti, B., Saykin, A.J., Sperling, R.A., Johnson, K.A., Schofield, P.R., Buckles, V., Morris, J.C., Benzinger, T.L.S., 2015. Partial volume correction in quantitative amyloid imaging. Neuroimage. https://doi.org/10.1016/j.neuroimage.2014.11.058

Su, Y., Flores, S., Wang, G., Hornbeck, R.C., Speidel, B., Joseph-Mathurin, N., Vlassenko, A.G., Gordon, B.A., Koeppe, R.A., Klunk, W.E., Jack, C.R., Farlow, M.R., Salloway, S., Snider, B.J., Berman, S.B., Roberson, E.D., Brosch, J., Jimenez-Velazques, I., van Dyck, C.H., Galasko, D., Yuan, S.H., Jayadev, S., Honig, L.S., Gauthier, S., Hsiung, G.Y.R., Masellis, M., Brooks, W.S., Fulham, M., Clarnette, R., Masters, C.L., Wallon, D., Hannequin, D., Dubois, B., Pariente, J., Sanchez-Valle, R., Mummery, C., Ringman, J.M., Bottlaender, M., Klein, G., Milosavljevic-Ristic, S., McDade, E., Xiong, C., Morris, J.C., Bateman, R.J., Benzinger, T.L.S., 2019. Comparison of Pittsburgh compound B and florbetapir in cross-sectional and longitudinal studies. Alzheimer’s Dement. Diagnosis, Assess. Dis. Monit. https://doi.org/10.1016/j.dadm.2018.12.008

Sun, D., van Erp, T.G.M., Thompson, P.M., Bearden, C.E., Daley, M., Kushan, L., Hardt, M.E., Nuechterlein, K.H., Toga, A.W., Cannon, T.D., 2009. Elucidating a Magnetic Resonance Imaging-Based Neuroanatomic Biomarker for Psychosis: Classification Analysis Using Probabilistic Brain Atlas and Machine Learning Algorithms. Biol. Psychiatry. https://doi.org/10.1016/j.biopsych.2009.07.019

Thompson, P.M., Gledd, J.N., Woods, R.P., MacDonald, D., Evans, A.C., Toga, A.W., 2000. Growth patterns in the developing brain detected by using continuum mechanical tensor maps. Nature. https://doi.org/10.1038/35004593

Thompson, Paul M., Hayashi, K.M., De Zubicaray, G.I., Janke, A.L., Rose, S.E., Semple, J., Hong, M.S., Herman, D.H., Gravano, D., Doddrell, D.M., Toga, A.W., 2004. Mapping hippocampal and ventricular change in Alzheimer disease. Neuroimage. https://doi.org/10.1016/j.neuroimage.2004.03.040

Thompson, Paul M, Hayashi, K.M., Sowell, E.R., Gogtay, N., Giedd, J.N., Rapoport, J.L., de Zubicaray, G.I., Janke, A.L., Rose, S.E., Semple, J., Doddrell, D.M., Wang, Y., van Erp, T.G.M., Cannon, T.D., Toga, A.W., 2004. Mapping cortical change in Alzheimer’s disease, brain development, and schizophrenia. Neuroimage 23, S2–S18. https://doi.org/10.1016/j.neuroimage.2004.07.071

Tosun, D., Chen, Y.-F., Yu, P., Sundell, K.L., Suhy, J., Siemers, E., Schwarz, A.J., Weiner, M.W., 2016. Amyloid status imputed from a multimodal classifier including structural MRI distinguishes progressors from nonprogressors in a mild Alzheimer’s disease clinical trial cohort. Alzheimer’s Dement. 12, 977–986. https://doi.org/10.1016/j.jalz.2016.03.009

Tosun, D., Joshi, S., Weiner, M.W., 2014. Multimodal MRI-based imputation of the A β + in early mild cognitive impairment. Ann. Clin. Transl. Neurol. 1, 160–170. https://doi.org/10.1002/acn3.40

Vanwinckelen, G., Blockeel, H., 2012. On estimating model accuracy with repeated cross-validation. 21st Belgian-Dutch Conf. Mach. Learn.

Vounou, M., Nichols, T.E., Montana, G., Initiative, A.D.N., others, 2010. Discovering genetic associations with high-dimensional neuroimaging phenotypes: a sparse reduced-rank regression approach. Neuroimage 53, 1147–1159.

Wang, Y., Chan, T.F., Toga, A.W., Thompson, P.M., 2009. Multivariate tensor-based brain anatomical surface morphometry via holomorphic one-forms, in: Lecture Notes in Computer Science (Including Subseries Lecture Notes in Artificial Intelligence and Lecture Notes in Bioinformatics). https://doi.org/10.1007/978-3-642-04268-3_42

Wang, Y., Lui, L.M., Gu, X., Hayashi, K.M., Chan, T.F., Toga, A.W., Thompson, P.M., Yau, S.T., 2007. Brain surface conformal parameterization using riemann surface structure. IEEE Trans. Med. Imaging. https://doi.org/10.1109/TMI.2007.895464

Wang, Y., Shi, J., Yin, X., Gu, X., Chan, T.F., Yau, S.T., Toga, A.W., Thompson, P.M., 2012. Brain surface conformal parameterization with the ricci flow. IEEE Trans. Med. Imaging. https://doi.org/10.1109/TMI.2011.2168233

Wang, Y., Song, Y., Rajagopalan, P., An, T., Liu, K., Chou, Y.Y., Gutman, B., Toga, A.W., Thompson, P.M., 2011. Surface-based TBM boosts power to detect disease effects on the brain: An N=804 ADNI study. Neuroimage. https://doi.org/10.1016/j.neuroimage.2011.03.040

Wang, Y., Yuan, L., Shi, J., Greve, A., Ye, J., Toga, A.W., Reiss, A.L., Thompson, P.M., 2013. Applying tensor-based morphometry to parametric surfaces can improve MRI-based disease diagnosis. Neuroimage. https://doi.org/10.1016/j.neuroimage.2013.02.011

Wang, Y., Zhang, J., Gutman, B., Chan, T.F., Becker, J.T., Aizenstein, H.J., Lopez, O.L., Tamburo, R.J., Toga, A.W., Thompson, P.M., 2010. Multivariate tensor-based morphometry on surfaces: Application to mapping ventricular abnormalities in HIV/AIDS. Neuroimage 49, 2141–2157. https://doi.org/10.1016/j.neuroimage.2009.10.086

Wu, J., Zhang, J., Shi, J., Chen, K., Caselli, R.J., Reiman, E.M., Wang, Y., 2018. Hippocampus morphometry study on pathology-confirmed Alzheimer’s disease patients with surface multivariate morphometry statistics, in: Proceedings - International Symposium on Biomedical Imaging. https://doi.org/10.1109/ISBI.2018.8363870

Yao, Z., Fu, Y., Wu, J., Zhang, W., Yu, Y., Zhang, Z., Wu, X., Wang, Y., Hu, B., 2018. Morphological changes in subregions of hippocampus and amygdala in major depressive disorder patients. Brain Imaging Behav. https://doi.org/10.1007/s11682-018-0003-1

Yin, W., Osher, S., Goldfarb, D., Darbon, J., 2008. Bregman Iterative Algorithms for $\ell_1$-Minimization with Applications to Compressed Sensing. SIAM J. Imaging Sci. 1, 143–168. https://doi.org/10.1137/070703983

Younes, L., Ratnanather, J.T., Brown, T., Aylward, E., Nopoulos, P., Johnson, H., Magnotta, V.A., Paulsen, J.S., Margolis, R.L., Albin, R.L., Miller, M.I., Ross, C.A., Investigators, P.-H., Coordinators of the Huntington Study, G., 2014. Regionally selective atrophy of subcortical structures in prodromal HD as revealed by statistical shape analysis. Hum Brain Mapp 35, 792–809. https://doi.org/10.1002/hbm.22214

Zhang, J., Fan, Y., Li, Q., Thompson, P.M., Ye, J., Wang, Y., 2017a. Empowering cortical thickness measures in clinical diagnosis of Alzheimer’s disease with spherical sparse coding, in: Proceedings - International Symposium on Biomedical Imaging. https://doi.org/10.1109/ISBI.2017.7950557

Zhang, J., Li, Q., Caselli, R.J., Thompson, P.M., Ye, J., Wang, Y., 2017b. Multi-source Multi-target Dictionary Learning for Prediction of Cognitive Decline. Springer, Cham, pp. 184–197. https://doi.org/10.1007/978-3-319-59050-9_15

Zhang, J., Shi, J., Stonnington, C., Li, Q., Gutman, B.A., Chen, K., Reiman, E.M., Caselli, R., Thompson, P.M., Ye, J., Wang, Y., 2016a. Hyperbolic space sparse coding with its application on prediction of Alzheimer’s disease in mild cognitive impairment, in: Lecture Notes in Computer Science (Including Subseries Lecture Notes in Artificial Intelligence and Lecture Notes in Bioinformatics). https://doi.org/10.1007/978-3-319-46720-7_38

Zhang, J., Stonnington, C., Li, Q., Shi, J., Bauer, R.J., Gutman, B.A., Chen, K., Reiman, E.M., Thompson, P.M., Ye, J., Wang, Y., 2016b. Applying sparse coding to surface multivariate tensor-based morphometry to predict future cognitive decline, in: Proceedings - International Symposium on Biomedical Imaging. https://doi.org/10.1109/ISBI.2016.7493350

Zhang, J., Tu, Y., Li, Q., Caselli, R.J., Thompson, P.M., Ye, J., Wang, Y., 2018. Multi-task sparse screening for predicting future clinical scores using longitudinal cortical thickness measures, in: Proceedings - International Symposium on Biomedical Imaging. https://doi.org/10.1109/ISBI.2018.8363835

